# A flexible Janus head: molecular determinants of a viral protein’s RNAi suppressor and capsid forming activities

**DOI:** 10.1101/2025.11.26.690650

**Authors:** Dennis Arendt, Selvaraj Tamilarasan, Ralph Peter Golbik, Iris Thondorf, Julian Bender, Hauke Lilie, Torsten Gursinsky, Christoph Parthier, Carla Schmidt, Milton T. Stubbs, Sven-Erik Behrens

## Abstract

Viral suppressors of RNA silencing (RNAi) expressed by plant viruses, VSRs, are exceptional proteins. Although not conserved, even within virus families, most VSRs bind small interfering RNAs, siRNAs, thereby blocking antiviral RNAi. *Turnip crinkle virus*, a member of the *Tombusviridae* family, encodes a VSR, TCV P38, which also forms the viral capsid. Biochemical studies of the purified protein revealed that the TCV P38 VSR functions as a metastable dimer that binds double-stranded (ds) RNA with high affinity *via* an induced-fit mechanism of both binding partners. Consistent with its role as a VSR that interferes with antiviral RNAi at various stages, P38 distinguishes between siRNAs of different lengths. Consistent with its capsid-forming function, the protein binds longer dsRNAs cooperatively. Structural data obtained from an RNA-free capsid-like icosahedral crystal and modeling of *Tombusviridae* capsid proteins suggest that flexible interactions between the P (protruding)-domains of P38 are important determinants for forming both the VSR dimer and the capsid structure. Studies with protein mutants confirmed this and also revealed the central role of the S (shell)- and R (RNA-binding)-domains of TCV P38 in adaptive substrate binding. Our study provides comprehensive insights into the molecular and structural properties of a versatile viral “Janus head” protein.

**Graphical abstract:** 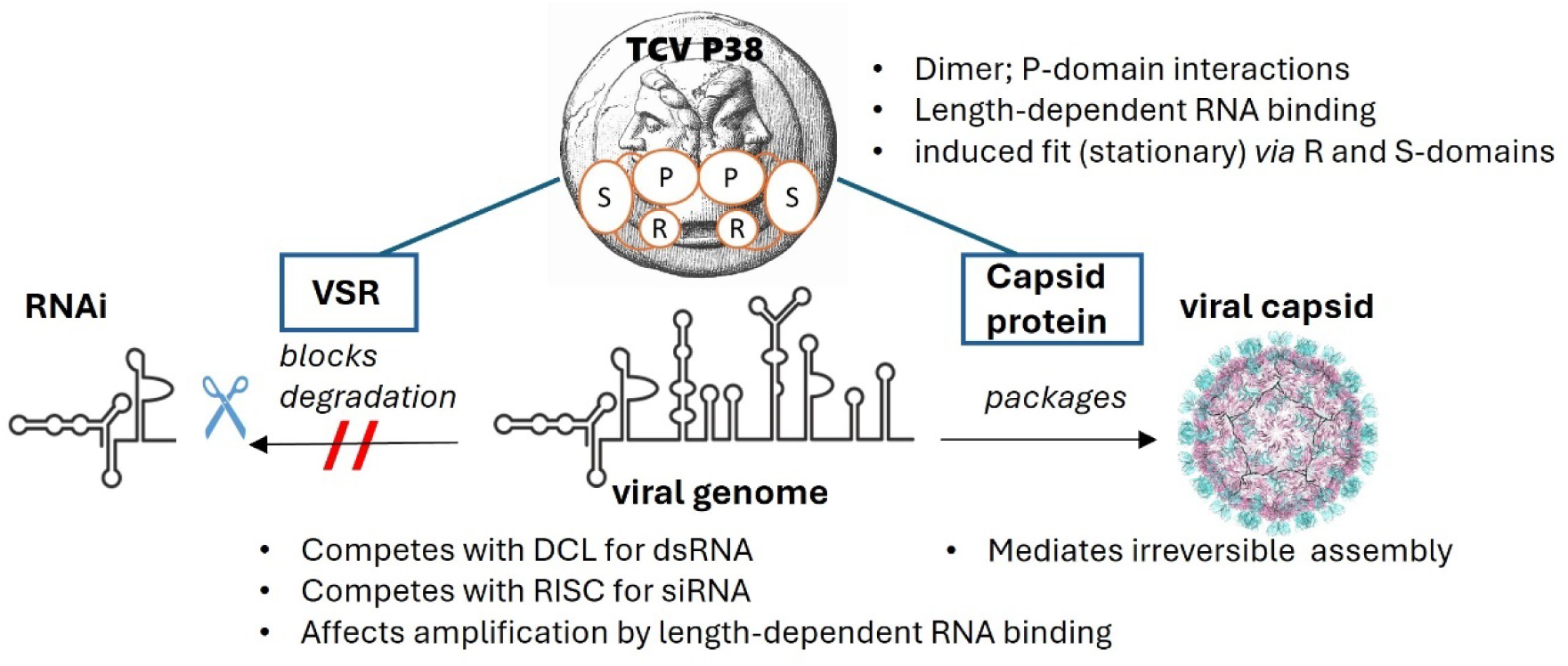

## Introduction

RNA silencing, also termed RNA interference (RNAi), is an evolutionarily conserved mechanism that regulates gene expression at the transcriptional or posttranscriptional level in eukaryotic cells (1,2). In addition to central functions in development and the maintenance of genome integrity, RNAi represents the primary immune system in plants for protection against viruses. RNAi is activated extracellularly by Ca^2+^-signaling (3–8) and intracellularly by double-stranded (ds) RNA acting as a pathogen-associated molecular pattern (PAMP): for example, in infections with positive-strand RNA viruses, which represent the majority of plant-infecting viruses, antiviral RNAi can be induced by viral dsRNA replication intermediates, structured regions of viral mRNAs and/or genomes, and dsRNAs that are generated from viral RNA templates by host RNA-dependent RNA polymerases (RDRs) (9,10).

Antiviral RNAi involves two endonuclease cleavage reactions (11,12). First, viral dsRNA PAMPs are recognized by the RNase III Dicer-like proteins DCL4 and DCL2 and processed into 21- and 22 nucleotides (nts)-long small interfering RNAs (siRNAs), respectively (13,14). RNase H-related Argonaute proteins, in plant antiviral processes mainly AGO1 and AGO2, then degrade the target RNA. The AGOs are part of RNA-induced silencing complexes (RISCs) and contain a bound siRNA guide strand that specifically interacts with the target RNA *via* base pairing (15–17). Antiviral RNAi is further enhanced by the biogenesis of secondary siRNAs. Together with the protein SGS3 (Suppressor of Gene Silencing 3), RDRs such as RDR6 convert single-stranded RNAs into new dsRNA substrates, which are then processed by DCLs (18–21). The accumulation and spread of primary and secondary (RDR-generated) siRNAs can lead to interference with viral protein synthesis and replication, a reduction in viral titer, and the development of temporary local and systemic plant immunity (**Supplementary Fig. S1**) (22,23).

RNA viruses, in particular, have high mutation rates and rapidly generate new quasispecies that evade protein-mediated immunity mechanisms, such as plant resistance R proteins (11,12,24). Antiviral RNAi can compensate for this as DCL recognition of viral dsRNA molecules is sequence independent, generating a broad repertoire (“pool”) of siRNAs. To counteract this, viruses have evolved suppressor proteins of RNA silencing (VSRs) that actively suppress antiviral RNAi at different stages (25–29): some VSRs inhibit siRNA biogenesis and accumulation, while others suppress the antiviral activity of siRNAs (9). Remarkably, VSRs encoded by different virus families often show no recognizable sequence similarity, suggesting independent origins (25,28). Most of the VSRs identified are also multifunctional. In addition to their role in suppressing RNA silencing, they can act as replicases, movement proteins, core (coat, capsid) proteins, components of viral transmission, proteases or regulators of transcription (30–33).

A remarkable example of a multifunctional VSR is the P38 protein of turnip crinkle virus (TCV; genus Betacarmovirus, family *Tombusviridae*), a positive-strand RNA virus with a genome encoding five proteins. On the one hand, TCV P38 encapsulates the viral genome. Like with many isometric viruses, the capsid is formed from 180 copies of P38 in an icosahedral T = 3 arrangement (**Supplementary Fig. S2**) (34,35). Each P38 monomer contains two consecutive jellyroll domains: a middle “shell” or S-domain (amino acids 80-242), and a C-terminal “protruding” or P-domain (amino acids 248-351). The N-terminal “RNA-binding” R-domain (amino acids 1-45), which does not follow the icosahedral shell symmetry and is therefore disordered in capsid structures, is suggested to serve as a binding site for the viral genome (35–38). Capsid formation is dispensable for virus replication, although active virion formation by P38 is required for long-distance movement, *i.e.*, virus exit from the vasculature and transport in systemically infected leaves (9,39–46). On the other hand, TCV P38 is an effective VSR (47). Using TCV mutants encoding an attenuated P38 VSR, the anti-TCV functions of DCL4, DCL2, the DCL4-supporting DRB4 protein, and AGO1 and AGO2 were revealed (39,48,49). There is evidence that P38 interferes with the activity of DCL2 and DCL4 and the processing of RDR6-generated dsRNA (48,50); other studies suggested P38 to interact with AGO1 *via* defined glycine-tryptophan (GW) motifs (51,52) and with SGS3 (53), thus affecting RISC activity and SGS3/RDR6-mediated production of secondary siRNAs, respectively. Finally, P38 was indicated to bind dsRNA and siRNA, preventing DCL-mediated processing into siRNAs and entering of siRNAs into RISC (summarized in **Supplementary Fig. S1**) (13,50).

We present here a comprehensive biochemical analysis of TCV P38 that reveals detailed insights into molecular features enabling the protein’s remarkable dual role in the viral life cycle.

## Materials and Methods

### Plasmid constructs

The plasmid pTCV containing the cDNA of the total genome of *Turnip crinkle virus* was a gift from Dr. Anne Simon (University of Maryland, USA). For expression, TCV P38-ORF was cloned *via* PCR into the pET-SUMO vector (Invitrogen). Mutations W26A, R57A, R74A, R84A and W274A were introduced by site-directed mutagenesis (see **Supplementary Table S1** for primers). Successful cloning and mutagenesis were verified via sequencing (Seqlab, Göttingen).

### Recombinant production and purification of ^14^N- and ^15^N-TCV P38 and variants

The fusion protein His^6^-SUMO-TCV P38 and variants thereof were recombinantly produced in *Escherichia coli* BL21-CodonPlus (DE)-RIPL. Protein purification was carried out as described in detail in the Supplementary data section (see scheme in **Fig. 1A**).

**Figure 1.**
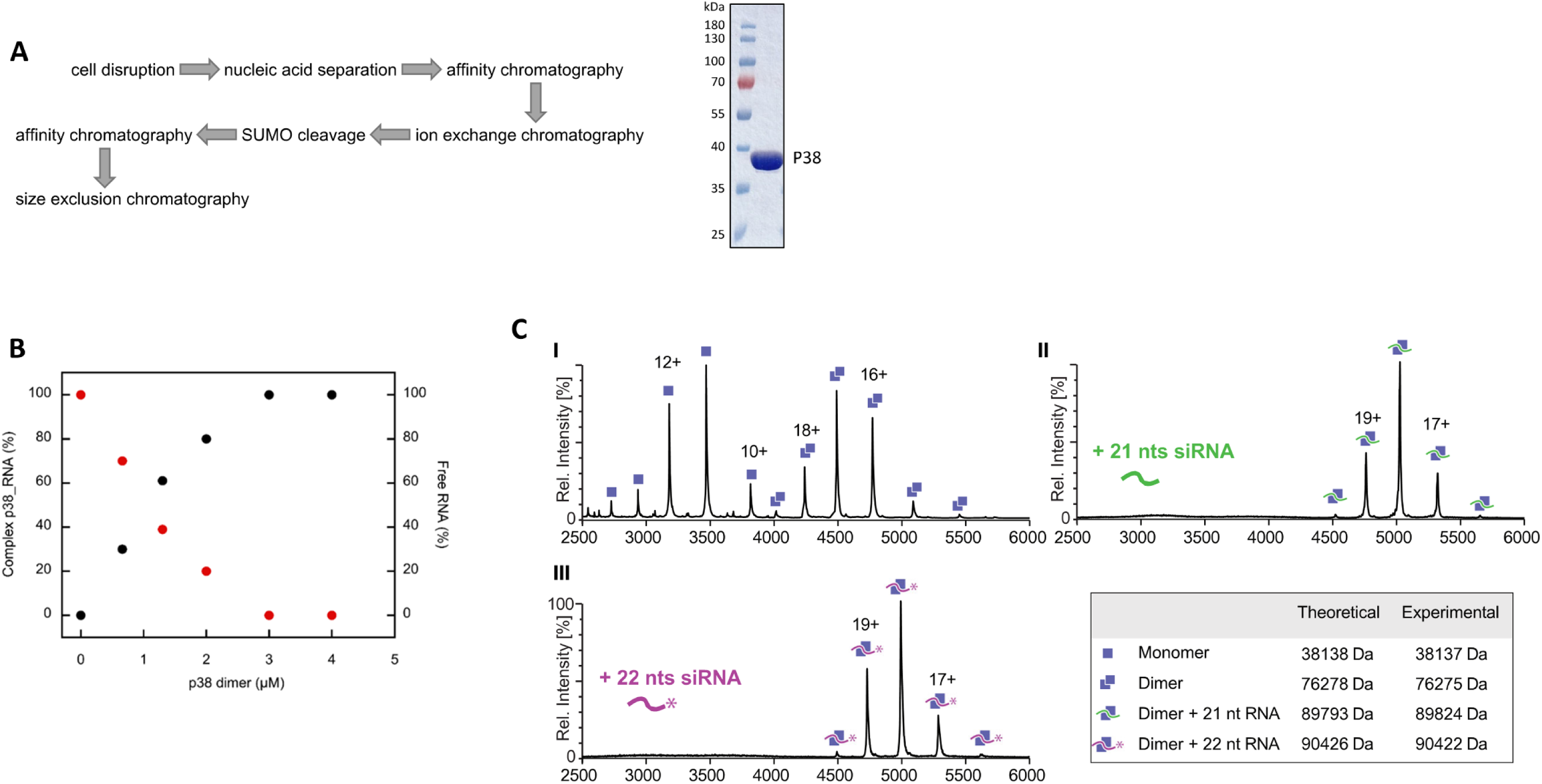
TCV P38 forms a dimer in solution. **(A)** (*left*) Purification workflow of TCV P38 and variants. Soluble proteins were purified according to the steps indicated, see Supplementary Materials and Methods for details. Inclusion bodies were isolated, solubilized and refolded as indicated in (75,76). (*right*) 4 µg of purified TCV P38 were analysed by Coomassie-stained SDS-PAGE. **(B)** Binding of TCV P38 to siR gf698_22 analysed by AUC (sedimentation velocity). Bound (black filled circles) and free (red filled circles) siRNA were detected by absorbance at 260 nm. Amplitudes of the respective signals were normalized. A binding stoichiometry of one siRNA molecule to one TCV P38 dimer was determined from titration. **(C)** Binding of TCV P38 to siRNA analysed by native MS. **(I)** Monomer (single squares) and dimer (double squares) peaks of TCV P38 demonstrate an equilibrium between the two species. Masses of charged states of the corresponding species are indicated. **(II** and **III)** Shift to a dimeric peak distribution on binding to siR gf698_21 (double squares with green line) and to siR gf698_22 (double squares with purple line). Theoretical and experimentally determined masses are indicated in the table.

### Analytical ultracentrifugation of TCV P38 and TCV P38_W274A

TCV P38 and TCV P38_W274A were analyzed at a protein concentration of 4 µM in 50 mM sodium phosphate, 0.1 M NaCl, pH 7.5 by an analytical ultracentrifuge (Optima XL-I BECKMAN) equipped with an An50Ti rotor and double sector cells. Sedimentation equilibrium measurements were carried out at 10,000 or 12,000 rpm, sedimentation velocity was monitored at 40,000 rpm at 20 °C every 10 min. The data was recorded at a wavelength of 230 nm and analyzed using the software Sedfit (54). In case of RNA binding, 2 µM siR gf698_22 was incubated with varying concentrations of TCV P38 in 50 mM TRIS, 100 mM NaCl, 1 mM TCEP, pH 7.6 and sedimentation velocity was monitored at 40,000 rpm. The amount of bound RNA was quantified at 260 nm by analyzing the amplitude of the rapidly sedimenting species (RNA bound to protein) and slowly sedimenting species (free RNA).

### Quantitation of RNA binding to TCV P38 and variants by electrophoretic mobility shift assay (EMSA)

The determination of dissociation constants (K_D_ values) of the recombinantly produced TCV P38 proteins and RNAs was performed by an EMSA approach (55). For direct measurements of the binding affinities, radiolabeled RNA (≤ 30 pM) was incubated with varying concentrations of the purified protein in binding buffer (20 mM TRIS, 100 mM NaCl, 1 mM EDTA, 1 mM DTT, 0.02 % Tween-20, pH 7.5) at 24 °C for 1-2 h. For competition experiments, the unlabeled but phosphorylated competitor RNA was titrated against a complex of radiolabeled siRNA (≤ 30 pM) that had been bound to a fixed concentration of 0.9 nM (dimer) TCV P38 in binding buffer overnight. The samples were mixed at 0.25 parts (v/v) with loading dye (50 mM TRIS pH 7.5, 10 mM EDTA, 0.002 % bromophenol blue, 0.002 % xylene cyanol, 50 % (v/v) glycerol) and analyzed by PAGE on a native 6 % TBE gel. The protein-bound and free RNAs were detected by phosphor-imaging (Storm 860, Molecular Dynamics) and quantified by ImageQuant software (GE Healthcare) (55). All measurements were performed in at least triplicate. With the exception of TCV P38_W274A, TCV P38 and variants are dimeric proteins, so that the normalized fractions of bound RNA were plotted against the total protein dimer concentrations and fitted to equation 1 for binding constant determination in direct binding reactions. Multiple binding (sigmoidal behavior) was considered in the cooperativity value.

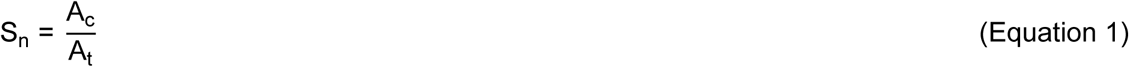

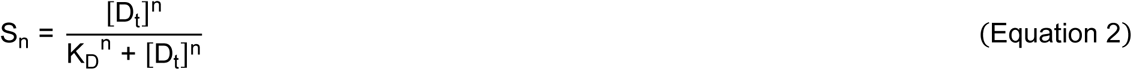

A_c_ is the radioactivity area of the complex in the PAGE lane and A_t_ the total radioactivity of the PAGE lane, respectively. S_n_ is the normalized fraction of bound RNA, [D_t_] the total protein dimer concentration, K_D_ the binding constant, and n the cooperativity. For the variant TCV P38_W274A, the total monomer concentration and a cooperativity of 2 were used. Parameters were additionally scrutinized by fitting the linearized data in reciprocal plots. For competition reactions, the fractions of bound radiolabeled siRNA were plotted against free competitor RNA concentrations and fitted to equation 3.

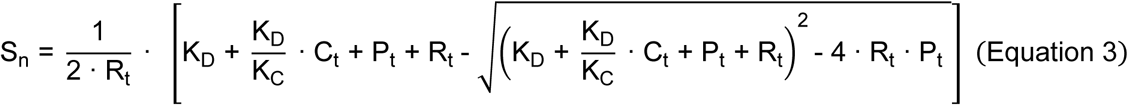

R_t_ is the total concentration of radiolabeled bound siRNA, C_t_ the total concentration of competitor RNA, P_t_ the total protein dimer concentration (here TCV P38), and K_C_ the apparent dissociation constant of the competitor RNA.

### Native mass spectrometry

^14^N- and ^15^N-TCV P38 in 50 mM sodium phosphate, 0.1 M NaCl, pH 7.5 were transferred into 200 mM ammonium acetate solution using 30K molecular weight cut-off filter devices (Vivaspin 500, Sartorius, Göttingen, Germany) and washing five times according to manufacturer’s protocols. Protein concentration after buffer exchange was determined by absorbance at 280 nm and the final protein concentration was adjusted to < 20 µM with ammonium acetate. Sample was loaded into in-house prepared gold-coated borosilicate capillaries (56) and analysed on a Waters Q-ToF Synapt G1 modified for the transmission of high masses. Typical analysis conditions were: capillary voltage, 1.3-1.7 kV; backing pressure, 4-8 mbar; sampling cone voltage, 120 V; extraction cone voltage, 5 V; collision cell pressure, 0.04 mbar; trap collision cell voltage, 10 V; transfer collision cell voltage, 10-30 V. Acquired mass spectra were processed with MassLynx (V4.1). For analysis of RNA-binding to TCV P38, the RNA was diluted into ammonium acetate to a concentration < 20 µM and directly added to the protein. After mixing, the sample was loaded into the capillary and analysed as described. For RNA removal, the sample was transferred into ammonium acetate buffer using centrifugal filters as described above.

### Crystallization and structure determination of TCV P38

The protein was concentrated to 8.8 mg·mL^-1^ (231 µM) using VIVASPIN concentrators (MWCO 50 kDa for dimers) in 50 mM TRIS, 100 mM NaCl, 1 mM TCEP, pH 7.6 and crystallized by hanging-drop vapour diffusion at 12 °C after mixing 2 µL protein solution with the same volume of reservoir solution. Crystals of TCV P38 grew within two weeks in 100 mM MES (sodium salt), 200 mM ammonium sulphate, 22 % (w/v) PEG 8000, pH 5.7. Diffraction data were collected under cryogenic conditions from a single crystal at the beamline BL 14.2 (BESSY II synchrotron, Helmholtz-Zentrum Berlin) using a hybrid photon counting detector (Pilatus3 2M, Dectris, Switzerland) and processed to 2.9 Å with the XDS software package (57) in the rhombohedral space group R3. Intensity statistics indicated merohedral twinning; complete data collection statistics are given in **Supplementary Table S2**. The structure was solved by molecular replacement employing the program PHASER (58) using coordinates of the *Carnation mottle virus* crystal structure (PDB entry: 1OPO, (59)) as search model. 20 non-crystallographic symmetry-related TCV P38 S-domain monomers could be located in the asymmetric unit of the crystal, forming four pentamers. The pseudo-capsid, consisting of 60 TCV P38 monomers with a triangulation number T = 1, is assembled by three asymmetric units of the crystal. Whereas each S-domain could be placed straightforwardly, initial electron densities for the P-domains were poor and could not be improved by non-crystallographic symmetry (NCS) averaging. Using phases from coordinates of the 20 S-domains only, a few β-strands each of the A and F P-domain monomers could be built. Taking these latter coordinates as a single rigid body, corresponding positions of the remaining nine P-domain dimers could be located, demonstrating the P-domain dimer as a conserved structural unit and determining directions of the local non-crystallographic twofold axes different to those of the S-domains. This in turn facilitated local NCS averaging of the ten P-domain dimers, allowing the entire P-domain to be traced. The final structure was completed by iterative model building using the program Coot (60) and NCS-restrained refinement (taking twinning into account) employing the Phenix software suite (61). No density was observed for the R-domains, probably due to non-adherence to the icosahedral NCS. Each of the 20 TCV P38 monomers is defined from residue D80 to I351, with each S-domain well defined. On the other hand, P-domains of the A and F monomers are best defined and C and D poorest. Structure validation was carried out using MolProbity (62), molecular figures were created with the software PyMOL (Schrödinger LLC, New York, USA). Refinement statistics are given in **Supplementary Table S2**.

### Database searches and model generation

To identify protein sequences that are homologous to the TCV P38 capsid protein, a BLAST (63) search was conducted using the sequences of the full-length TCV P38, the R-domain (amino acids 1-45), the S- domain (amino acids 82-242), and the P-domain (amino acids 248-351) in the UniProtKB reference proteomes and Swiss-Prot databases (64). The BLOSUM62 matrix (65) and the default parameters for the handling of gaps were utilized. Protein sequences of different strains of the same virus and of unclassified RNA viruses were manually excluded from the dataset. This process yielded a total of 36 homologous sequences. The sequences were then submitted to the AlphaFold 3 server (66,67) which generated structural models of the coat proteins. Visual inspection of the models revealed that the two species assigned to the Gammacarmovirus genus, *Cowpea mosaic virus* (CPMoV) and *Soybean yellow mottle mosaic virus* (SYMMV), exhibited a significant deviation from all other capsid proteins with regard to the structure of the P-domain (data not shown). As a result, these two species were excluded from the subsequent stages of the investigation. The initial alignment of all models was conducted using the sequence and structural alignment option within MOE (68). This was followed by the superposition of residues corresponding to the respective domain. Subsequently, dendrograms were constructed based on the RMSD values in MOE. The sequence and structure alignment scheme for each domain was prepared with ESPript (69) and illustrations of the AF3-generated models were prepared with MOE (70).

### Cell culture and preparation of BYL

*Nicotiana tabacum* BY-2 cells were cultured at 23 °C in Murashige-Skoog liquid medium (Duchefa Biochemie, Haarlem, Netherlands). Evacuolated BY-2 protoplasts were obtained by percoll gradient centrifugation and cytoplasmic extract (BYL) prepared as described earlier (71,72).

### Dicer cleavage and inhibition assay. *In vitro* Slicer assay

The BYL extract displays an intrinsic DCL (dicer) activity. To assay this enzymatic reaction, 12.5 nM double-stranded TCV 3’UTR labelled with [α-^32^P] CTP was used as target. The substrate was simultaneously incubated with TCV P38 proteins at a concentration of 100 nM in 50 % (v/v) BYL at 25 °C for 2.5 h under previously described conditions (73,74). Total RNA was isolated from the reaction by treatment with 20 µg proteinase K in the presence of 0.5 % SDS for 30 min at 37 °C, followed by chloroform extraction and ethanol precipitation. ^32^P-labeled products were separated on 12.5 % TBE polyacrylamide gels containing 8 M urea and visualized by phosphor-imaging (Storm 860, Molecular Dynamics). In addition, the Dicer assay was performed in a pre-incubation manner to allow complex formation between TCV P38 proteins and RNA (30 min) before application to dicer activity. To generate siRNA-programmed AGO1/RISC *in vitro*, *NbAGO1-1L* mRNA was *in vitro* translated in 50 % (v/v) BYL under the previously described conditions (73,74). Briefly, 0.5 pmol *AGO1* mRNA was translated in a 20 µl reaction in the presence of 10 nM siR gf698_21 duplex. For slicer assays in the presence of VSR, 100 nM purified VSR was applied to the reaction at the same time point as the siRNA and the *AGO1* mRNA (simultaneous incubation). Finally, the slicer assay was performed in a post-incubation manner with a delayed application (90 min) of the respective VSR. After a total incubation time of 2.5 h at 25 °C, 2 µg of firefly luciferase (competitor) mRNA and the ^32^P-labeled target RNA (50 fmol) were added. The cleavage reaction was performed for 15 min. Total RNA was isolated from the reaction by treatment with 20 µg proteinase K in the presence of 0.5 % SDS for 30 min at 37 °C, followed by chloroform extraction and ethanol precipitation. ^32^P-labeled products were separated on 5 % TBE polyacrylamide gels containing 8 M urea and visualized by phosphor-imaging (Storm 860, Molecular Dynamics).

### Further information on Materials and Methods can be found in the Supplementary data section

It covers the following topics: Recombinant production and purification of ^14^N- and ^15^N-TCV P38 and variants. Origin, generation and modification of RNAs. Circular dichroism studies of TCV P38 and variants.

## Results

### Purification and oligomerization behavior of TCV P38

We produced TCV P38 recombinantly in *E. coli* as a soluble fusion protein. Following several purification steps and cleavage/separation of the His_6_-SUMO fusion/tag-unit used (**Fig. 1A**), the protein was obtained in homogeneous form with authentic termini and free of any nucleic acids. This was verified by tryptic digestion and LC-MS/MS analysis, as well as intact mass using native mass spectrometry. Importantly, analytical ultracentrifugation (AUC) revealed that the purified TCV P38 protein predominantly forms a dimeric species, both in the absence and in the presence of a bound siRNA. In the latter case, we determined a stoichiometry of one molecule of TCV P38 protein dimer to bind one siRNA molecule (**Fig. 1B** and **C** and **Supplementary Fig. S3**). Dimer formation was confirmed using native MS (**Fig. 1C**; see Materials and Methods). Interestingly, native MS in the absence of RNA showed the presence of monomeric and dimeric P38 species suggesting that the RNA-free dimer is less stable than the RNA-bound form (**Fig. 1C**).

### TCV P38 binds a variety of dsRNAs with similar affinity

The AUC and native MS data of the purified protein confirmed earlier observations (50) that TCV P38 binds dsRNA with high affinity (**Fig. 1**). Here, quantitative measurements on the purified protein allow a detailed analysis of this property. We applied an earlier established direct electrophoretic mobility shift assay (55) to determine the dissociation constants (K_D_ values) of TCV P38 for various species of RNA substrates, namely 21, 22 and 24 nts variants of the siRNA siR gf698 (directed against the mRNA of green fluorescent protein), the microRNAs miR162, miR168 and miR403 from *Arabidopsis thaliana* (*At*) and *Nicotiana benthamiana* (*Nb)* (55), a synthetic 33 nts dsRNA (dsRNA_33) with random nucleotide composition (77) and the *in vitro* transcribed 3’ untranslated region of the TCV RNA genome, TCV 3’UTR. The TCV 3’UTR is known to be highly structured and to contain defined dsRNA elements (78).

These experiments showed similar, very high binding affinities for each of the RNAs tested, with K_D_ values in the pico- to nanomolar range (**Table 1**; **Fig. 2A**). Importantly, the EMSA results demonstrated that TCV P38 forms one complex each with the siRNAs and miRs, but two or multiple complexes with the dsRNA_33 and the viral 3’UTR, respectively (**Fig. 2A** and **2B**; **Supplementary Fig. S4**). Fitting the binding isotherms to the data from the “longer” RNAs yielded similar K_D_ values as with the siRNAs, but with a cooperativity value of 2, indicating a stepwise complex formation with multiple independent binding events (**Table 1**; **Fig. 2C**). In contrast, TCV P38 has little to no affinity for truly single-stranded RNAs such as individual siRNA guide strands (**Table 1**).

**Figure 2.**
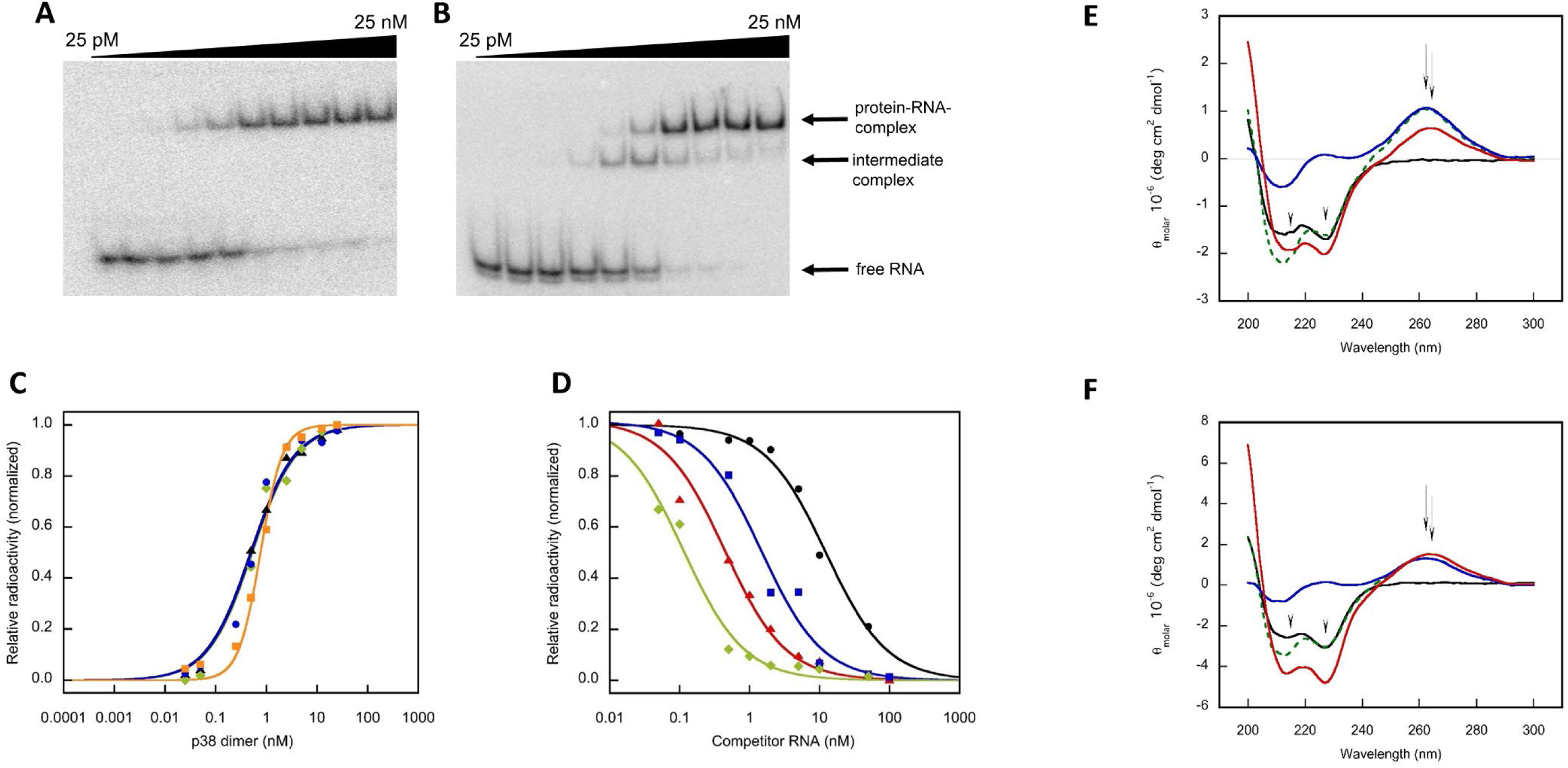
RNA binding by TCV P38 monitored by EMSA. The protein dimer concentration is indicated on top of the gels **(A** and **B)**. Free and bound (complexed) RNAs were detected in the analytical gel. **(A)** Binding of TCV P38 to siR gf698_21. **(B)** Binding of TCV P38 to dsRNA_33. **(C)** Binding isotherms of TCV P38 to siR gf698_21 (black triangle), siR gf698_22 (green diamond), siR gf698_24 (blue circle) and dsRNA_33 (orange square). The sigmoidal binding to the longer dsRNA revealed a cooperativity value of 2 in the fitting routine (**Table 1**). **(D)** Competitive binding of different RNAs on a preformed complex of TCV P38 and radioactively labelled siRNA was monitored by EMSA. Releasing isotherms of the original siRNA are displayed for competitor RNAs of varying lengths, such as siR gf698_21 (black circle), siR gf698_22 (red triangle), siR gf698_24 (blue square) and dsRNA_33 (green diamond). Data fitting considering the known K_D_ value of the preformed complex yielded K_C_ values of the competitor RNAs that reflect their ability to replace the radioactively labelled siRNA in an equilibrium reaction. The approach was performed for different combinations of original siRNAs and competitor RNAs (**Table 2**). Complex formation of TCV P38 and siR gf698_21 **(E)** as well as siR gf698_24 **(F)** was analyzed by far and near UV circular dichroism. Spectra of TCV P38 (black line), siR gf698 (blue line), and TCV P38 complexed with siRNA (red line) are shown. The spectrum of TCV P38 displayed α-helical secondary structure elements with a negative extremum at 227 nm (indicated by arrow). A larger signal amplitude indicates an increase in α-helical secondary structure of the protein during complex formation. The siRNAs displayed a spectrum with a positive extremum at 260 nm (indicated by arrow). A slight bathochromic shift to 262 nm (indicated by arrow) was detected during complex formation, which is accompanied by a change in the signal amplitude. Note that the additive spectrum of the individual components (green dashed line) does not match the spectrum of the complex. The observed structural changes of both components reveal an induced fit in complex formation.

**Table 1.** Complex formation of TCV P38 wildtype protein and different RNA molecules.

| RNA | $K_D$ (nM) | cooperativity $n^1$ |
| --- | --- | --- |
| siR gf698_21 | $0.56 \pm 0.09$ | 1 |
| siR gf698_22 | $0.53 \pm 0.07$ | 1 |
| siR gf698_24 | $0.44 \pm 0.09$ | 1 |
| dsRNA_33 | $0.77 \pm 0.05$ | 2 |
| AtmiR162 | $0.30 \pm 0.03$ | 1 |
| AtmiR168 | $0.55 \pm 0.05$ | 1 |
| AtmiR403a | $0.87 \pm 0.13$ | 1 |
| NbmiR403a | $0.36 \pm 0.05$ | 1 |
| NbmiR403b | $1.27 \pm 0.10$ | 1 |
| ss gf698_21 | n.a. | n.a. |
| ss gf698_24 | n.a. | n.a. |
| TCV 3'UTR | $0.68 \pm 0.08$ | 2 |

<sup>1</sup> Cooperativity denotes the number of dimers that bind to one molecule RNA.
Data fitting and investigation was performed on the basis of the dimer concentration.
n.a. not applicable - no binding detectable
siR – siRNA with length of 21, 22 and 24 nts, respectively
miR – microRNA from *Arabidopsis thaliana* (At) or *Nicotiana benthamiana* (Nb) with lengths of 21 and 22 nts, respectively (55)
dsRNA – synthetic double-stranded RNA with a length of 33 nts of random sequence
ss – single stranded (guide strand) of respective siRNA
gf698 – GFP mRNA target (50)

**Table 2.** Competitive binding constants of the TCV P38 wildtype protein to different RNAs determined by electrophoretic mobility shift assay (EMSA)

| RNA (radioactively labelled) for complex formation | RNA (unlabelled) competitor | $K_C$ (nM) | $K_{rel} = \frac{K_C}{K_D}$ |
| --- | --- | --- | --- |
| siR gf698_21 | siR gf698_21 | $4.50 \pm 1.20$ | $8.82 \pm 2.38$ |
| siR gf698_21 | siR gf698_22 | $0.18 \pm 0.05$ | $0.34 \pm 0.10$ |
| siR gf698_21 | siR gf698_24 | $0.40 \pm 0.10$ | $0.82 \pm 0.20$ |
| siR gf698_21 | dsRNA_33 | $0.14 \pm 0.03$ | $0.23 \pm 0.05$ |
| siR gf698_21 | AtmiR162 | not replaceable at concentrations < 25 nM of the competitor RNA |  |
| siR gf698_21 | AtmiR168 | not replaceable at concentrations < 25 nM of the competitor RNA |  |
| siR gf698_22 | siR gf698_21 | $0.70 \pm 0.07$ | $1.37 \pm 0.15$ |
| siR gf698_22 | siR gf698_22 | $0.57 \pm 0.06$ | $1.08 \pm 0.12$ |
| siR gf698_22 | siR gf698_24 | $0.33 \pm 0.03$ | $0.67 \pm 0.06$ |
| siR gf698_22 | dsRNA_33 | $0.07 \pm 0.03$ | $0.12 \pm 0.05$ |
| siR gf698_22 | AtmiR162 | $0.50 \pm 0.15$ | $1.35 \pm 0.41$ |
| siR gf698_22 | AtmiR168 | $0.95 \pm 0.30$ | $2.32 \pm 0.74$ |
| siR gf698_24 | siR gf698_21 | $1.40 \pm 0.16$ | $2.75 \pm 0.33$ |
| siR gf698_24 | siR gf698_22 | $0.35 \pm 0.10$ | $0.66 \pm 0.19$ |
| siR gf698_24 | siR gf698_24 | $0.33 \pm 0.08$ | $0.67 \pm 0.16$ |
| siR gf698_24 | dsRNA_33 | $0.06 \pm 0.01$ | $0.10 \pm 0.02$ |
| siR gf698_24 | AtmiR162 | $0.33 \pm 0.17$ | $0.89 \pm 0.46$ |
| siR gf698_24 | AtmiR168 | $0.70 \pm 0.20$ | $1.71 \pm 0.49$ |

Abbreviations, see **Table 1**.
Data fitting and investigation were performed on the basis of the dimer concentration.

Taken together, this data suggests that one TCV P38 dimer binds with high affinity to approximately 13 to 15 nt of dsRNA. Longer RNA stretches (>∼24 nt) apparently recruit additional copies of the dimers, which may correspond to the process of TCV P38 capsid formation (see Discussion).

### TCV P38 discriminates between different RNA substrates

When acting as a VSR, the key function of TCV P38 is assumed to be the effective sequestration of potentially antiviral siRNAs produced by plant DCLs immediately after infection and/or during a systemic RNAi response. In this context, it was of great interest to understand how well the protein discriminates between dsRNAs of different lengths. For this purpose, we established a competition assay. First, the TCV P38 dimer was saturated with radiolabeled 21, 22, or 24 nts siRNAs according to the measured K_D_ values (**Table 1**) to achieve a protein saturation rate of 60-80%, a value at which competition experiments could be easily performed. The formed complexes were then exposed to increasing concentrations of unlabeled competitor siRNAs of different sizes (**Fig. 2D**). Data was plotted to yield a binding isotherm and analyzed according to a single competitor model from which the respective competitor constant K_C_ (the apparent dissociation constant of the competitor RNA) could be determined (**Table 2**) (55).

Unexpectedly, we observed that the siR gf698_21 in complex with the TCV P38 protein was barely outcompeted by the same 21 nts siRNA or the similarly long *At*miR162 or *At*miR168. However, siR gf698_21 could be replaced by siR gf698_22, siR gf698_24 or dsRNA_33 with K_C_ values of 0.18 nM, 0.40 nM and 0.14 nM, respectively (**Table 2**). When the same approach was performed starting from complexes of TCV P38 and siRs gf698_22 and 24 and competing siRNAs of different sizes (21, 22 and 24 nts), we observed that siR gf698_21 replaced/outcompeted the corresponding siR gf698_22 (K_C_ = 0.70 nM) and siR gf698_24 (K_C_ = 1.40 nM), but the latter only with low efficiency. In contrast, siR gf698_22 and siR gf698_24 was able to efficiently replace each other (K_C_ = 0.57 nM and 0.33 nM or 0.35 nM and 0.33 nM, respectively) (**Table 2**). In all combinations, dsRNA_33 turned out to be the most effective competitor (K_C_ ranges from 0.06 nM to 0.14 nM) (**Table 2**). This data is supported by the fact that comparable values were obtained in TCV P38 competition experiments for siR gf698_21 versus viral siRNAs of 21 and 22 nts derived from the *Tomato bushy stunt virus* genome (TBSV) (79) (not shown).

Our observations demonstrate that TCV P38 can discriminate between dsRNAs of different lengths, but not between those of different sequence composition. In simple terms, the longer the dsRNA, the better it is bound by the viral VSR.

### Co-folding/induced fit of TCV P38 and binding siRNA

The different K_C_ values measured in the competition experiments with different dsRNA setups (**Table 2**) indicate divergent conformational changes of TCV P38 upon binding of the different dsRNA ligands. To investigate this in further detail, we applied circular dichroism in far and near UV of protein-RNA complexes of TCV P38 100% saturated with either siR gf698_21, 22 or 24. Based on the EMSA data and the determined K_D_ values (**Table 1**), no free ligands were present in solutions that contained equimolar concentrations of the interaction partners. The spectra of the two individual components and of the complex were measured and superimposed. As shown in **Fig. 2E** and **Fig. 2F**, the theoretical sum of the spectra of the individual components diverged from the actual measured spectrum of the complex. Moreover, subtracting the spectrum of the siRNA alone from the spectrum of the TCV P38/siRNA complex and comparing the difference spectrum with that of TCV P38 alone highlighted differences in secondary structure elements. Interestingly, this was observed for both the protein (large amplitude change in far-UV circular dichroism – signals related to α-helical secondary structure elements) and the bound RNA (small amplitude and maximum change in near-UV circular dichroism – signals related to the nucleobases). The increase in amplitude of the α-helical part of the protein was directly proportional to the length of the bound siRNA. Accordingly, the data obtained with siR gf698_21 (**Fig. 2E**) and siR gf698_22 (not shown) differed marginally, while the conformational changes were pronounced in the complex containing the siR gf698_24 (**Fig. 2F**; see also below).

In sum, the obtained dataset suggests an induced fit mechanism due to co-folding of the protein and RNA binding partners during complex formation. As shown further below, the jellyroll-like folds of the S- and P-domains of TCV P38 exhibit a measurable but limited structural flexibility. Therefore, it is reasonable to assume that the adaptation process is primarily driven by structural stabilization and conformational changes in the R-domain of the protein (see Discussion section).

### TCV P38 is an interconverting dimer with rapid monomer exchange, a property that is blocked by the binding of RNA

To obtain further information on the characteristics of the induced fit of the TCV P38 upon RNA binding, we once more used native MS. For this purpose, we recombinantly expressed the protein in the presence of ^14^N- or ^15^N-nitrogen and purified the corresponding isotope-labeled variants. The mass shift introduced by the heavy isotope allowed to directly distinguish both populations by LC-MS/MS based on total mass. To confirm protein integrity, ^15^N-TCV P38 was first analyzed in the absence of RNA and confirmed to have the same dimer peak and charge state distribution as the ^14^N-TCV P38 form (**Fig. 3A**). Interestingly, when both forms were mixed at equimolar ratio to test for the formation of mixed dimers between the two populations (see **Supplementary Table S3** for exact conditions), we observed a characteristic peak distribution with a 1:2:1 intensity ratio corresponding in mass to a ^14^N/^14^N dimer, a mixed ^14^N/^15^N dimer, and a ^15^N/^15^N dimer, respectively (**Fig. 3A**). As both forms were mixed immediately prior to the MS analysis, the experiment indicated a rapid (< 5 min) kinetically controlled exchange of monomers between the differently labeled dimers. In close correlation with the previous AUC and MS data (**Fig. 1**), these observations highlight the metastable nature of the TCV P38 dimer.

**Figure 3.**
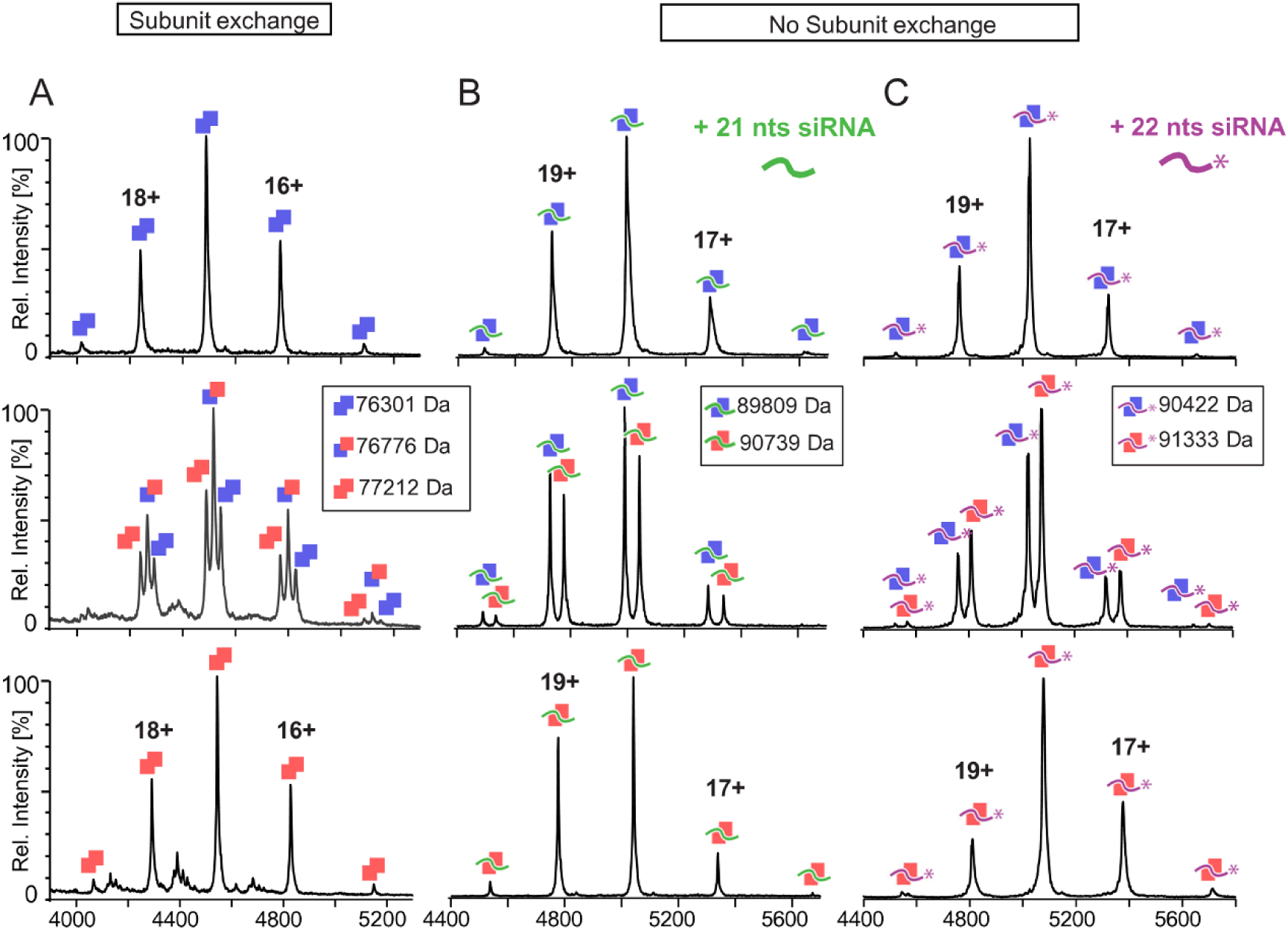
RNA binding by TCV P38 blocks subunit (monomer) exchange in the dimers. **(A)** Native mass spectrometry performed with TCV ^14^N-P38 (*top*, blue) and TCV ^15^N-P38 (*bottom*, red) revealed a similar dimer peak distribution (double squares). Charge states of selected peaks are indicated. An equimolar mixture of TCV ^14^N-P38 and TCV ^15^N-P38 (*middle*) revealed a 1:2:1 intensity distribution of the isotopically labeled monomers within the dimers, explained by subunit exchange between dimers. **(B)** Complex formation of TCV ^14^N-P38 (*top*) or TCV ^15^N-P38 (*bottom*) and siR gf698_21 resulted in the formation of stable dimers. The peak series correlated with the masses of the respective dimers with bound siRNA (blue or red double squares with green line). An equimolar mixture of the two complexes yielded a peak series corresponding to a superposition of the two separate spectra (*middle*), indicating that preformed dimers with bound siRNA do not exchange. **(C)** Experiments using siR gf698_22 revealed corresponding results (blue or red double squares with purple line) to those described in **(B)**. Masses determined are indicated in the inserts.

Next, we repeated the experiment in the presence of the siR gf698_21 and 22 (see **Supplementary Table S3**): *i.e.*, the ^14^N- and ^15^N-labeled forms of TCV P38 were pre-incubated with the RNAs and subsequently mixed. In accordance with our earlier data (**Fig. 1**), only RNA-bound protein dimers were detected and no peaks for TCV P38 without the nucleic acid were detected (top and bottom spectra). However, in contrast to the mixing experiment with the RNA-free protein isotope forms, no ^14^N/^15^N dimers could be detected that contained bound RNA (**Fig. 3B** and **3C**, middle parts). This indicated that the protein dimers remained stably associated after capture of the dsRNA ligand and that intermolecular monomer-subunit exchange between the dimers was abolished or greatly reduced compared to the RNA-free protein form.

Based on this data, we conclude that conformational immobilization - *i.e.* the transition from a “quasi-stationary” to a stable “stationary” dimer - is an essential functional component of the binding mechanism of dsRNA by TCV P38.

### GW motifs determine the structure and RNA binding capabilities of TCV P38

As already outlined, two GW-motifs (a highly conserved G25_W26 in the R-domain and a unique, TCV-specific G273_W274 in the P-domain) have previously been shown to be essential to the VSR function of TCV P38. To determine the role of these motifs in the structure and RNA binding function of TCV P38, three previously described protein variants (51,52), TCV P38_W26A, TCV P38_W274A and TCV P38_W26A_W274A, were generated, in which the GW motif tryptophans were replaced by alanines. Conveniently, the mutant proteins could be purified using the same procedure as the wildtype protein (**Fig. 1**).

Far UV circular dichroism measurements revealed considerable differences between the mutants: TCV P38_W26A_W274A was found to be completely unstructured, and TCV P38_W26A showed a strong decrease in the amplitude of the α-helical signal, indicating a strong perturbation of the helicity and thus the structural integrity of the protein. In contrast, TCV P38_W274A showed similar dichroic properties to TCV P38, but with a slightly reduced signal amplitude (**Fig. 4A**). The RNA binding properties of the mutant proteins, as measured by EMSA, were consistent: the double mutant protein exhibited virtually no RNA binding, and RNA binding was inhibited in the single GW mutants (**Table 3**).

**Figure 4.**
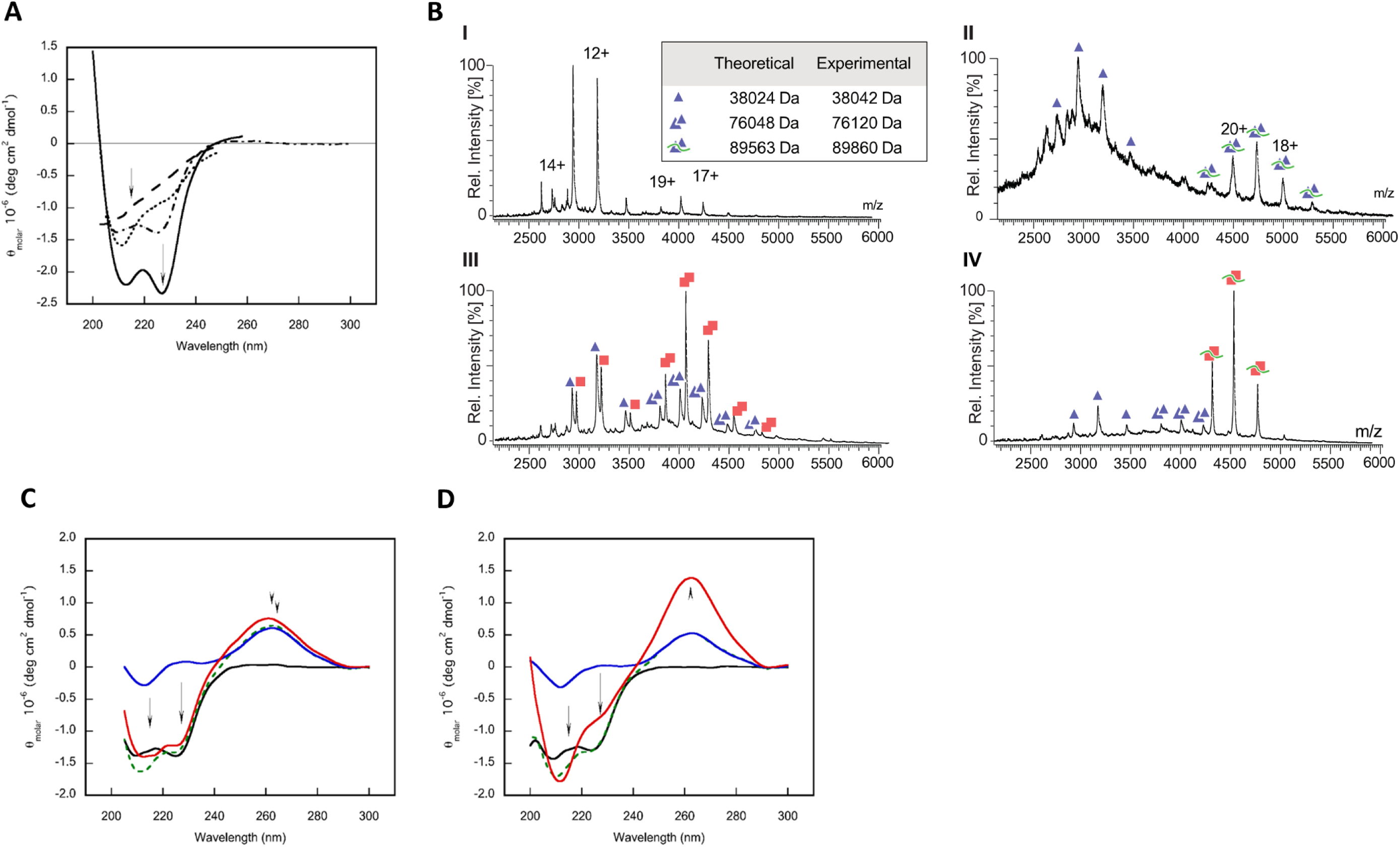
RNA binding of the TCV P38 tryptophan variants. **(A)** Far and near UV circular dichroism of TCV P38 and tryptophan variants. The spectrum of the wildtype protein is shown as black solid line (extrema marked by arrows). TCV P38_W26A_W274A displayed the spectrum of an unstructured protein (black dashed line). The spectrum of TCV P38_W26A (black dotted line) revealed half the amplitude of the wildtype protein and no pronounced extremum at 227 nm, indicating a strong perturbation of helicity. The spectrum of TCV P38_W274A (black dot-dashed line) displayed a similar form to the wildtype protein. However, a lower signal amplitude (arrows) indicates some structural perturbation. **(B)** Binding of TCV P38_W274A to siRNA analysed by native MS (nomenclature as in Fig. 3). **(I)** Monomers and dimers of the variant revealed an equilibrium in the absence of RNA; charge states of selected peaks are indicated. The peak series of the monomer was of higher intensity than that of the dimer implying a lower propensity to form oligomers. **(II)** Addition of siR gf698_21 resulted in complex formation with TCV P38_W274A as dimers (double triangles with green line). Free monomeric protein is indicated (single triangles). **(III)** Equimolar addition of TCV ^15^N-P38 to TCV ^14^N-P38_W274A did not result in the formation of mixed dimers. The peak distributions corresponded to monomers and dimers of either the isotopically labelled wildtype (single or double red squares) or TCV ^14^N-P38_W274A (single or double blue triangles). **(IV)** Equimolar addition of siR gf698_21 at the concentration related to a single protein component of the composition obtained from **(III)** shifted the TCV ^15^N-P38 entirely to the complex form (red double squares with green lines). No complexes were detected for TCV ^14^N-P38_W274A. Theoretical and experimentally determined masses are indicated in the table. **(C)** TCV P38_W274A shows an impaired binding of dsRNA in comparison to the wildtype (see also Fig. 2F and **Table 3**). Complex formation was performed at equimolar concentrations of siR gf698_21 considering the protein as dimer. Upon binding in cationic buffer (50 mM TRIS, 100 mM NaCl, 1 mM TCEP, pH 7.6) the circular dichroism spectrum of the complex (red solid line) displayed a slightly lower ellipticity at 227 nm compared to the monomer (black solid line). The spectral contribution of the dsRNA in the complex displayed both a slight hypsochromic shift and a slight increase in the ellipticity at 260 nm compared to the free ligand (blue solid line). **(D)** Performing the experiment in anionic buffer (50 mM sodium phosphate, 100 mM NaCl, 1 mM TCEP, pH 7.6) circular dichroism spectrum of the complex (red solid line) displayed lower ellipticity at 227 nm and higher ellipticity at 210 nm compared to the monomer (black solid line). The spectral region corresponding to the dsRNA displayed a twofold increase of the signal amplitude compared to the free ligand (blue solid line). Note that the additive spectra of the individual components (green dashed line) do not match the spectra of the respective complexes. The distinctive changes of the chirooptical properties of the complexes in comparison to the single components reveal induced conformational changes of both ligands.

**Table 3.** Complex formation of TCV P38 wildtype and mutant proteins and different RNA molecules.

| variant | RNA | K<sub>D</sub> (nM) | cooperativity n <sup>1</sup> |
| --- | --- | --- | --- |
| wildtype | siR gf698_21 | 0.56 ± 0.09 | 1 |
| wildtype | dsRNA_33 | 0.77 ± 0.05 | 2 |
| W26A | siR gf698_21 | 23.29 ± 1.60 | 1 |
| W274A <sup>2</sup> | siR gf698_21 | 42.77 ± 3.70 | 2 <sup>4</sup> |
| W274A <sup>3</sup> | siR gf698_21 | 517.88 ± 42.13 | 2 <sup>4</sup> |
| W26A_W274A | siR gf698_21 | > 100 | n.d. |
| R57A | siR gf698_21 | 4.09 ± 0.67 | 1 |
| R57A | dsRNA_33 | 1.09 ± 0.08 | 2 |
| R74A | siR gf698_21 | 0.36 ± 0.04 | 1 |
| R74A | dsRNA_33 | 0.22 ± 0.06 | 2 |
| R84A | siR gf698_21 | 7.17 ± 2.57 | 1 |
| R84A | dsRNA_33 | 10.46 ± 0.80 | 2 |

<sup>1</sup> Cooperativity denotes the number of dimers that bind to one molecule RNA.
Data fitting and investigation was performed on the basis of the dimer concentration.
<sup>2</sup> Binding experiments performed in 50 mM TRIS buffer, pH 7.6.
<sup>3</sup> Binding experiments performed in 50 mM sodium phosphate buffer, pH 7.6.
<sup>4</sup> Originally, the variant TCV P38\_W274A is a monomer and forms dimers upon binding dsRNA in a cooperative manner (**Fig. 4**). Data fitting and investigation was performed on the basis of the monomer concentration.
n.d. not determined

The TCV P38_W274A variant was further characterized by AUC and native MS in the absence and presence of bound dsRNA (**Fig. 4B** and **4C**; **Supplementary Fig. S5**). Interestingly, the data revealed that this mutant protein exists preferentially as a monomer in the absence of bound RNA (**Fig. 4B**). However, in the presence of the RNA ligand, TCV P38_W274A forms dimers, although the nucleic acid-induced dimer formation was less pronounced than for TCV P38 (**Fig. 4B**). Testing for the formation of mixed wildtype and mutant protein dimers using ^15^N-labeled forms (see **Supplementary Table S3** for conditions), showed that hybrid dimer formation between TCV P38 and TCV P38_W274A does not occur. In the presence of RNA, we observed that the nucleic acid was preferentially bound by the wildtype protein according to the lower K_D_ value and again no mixed dimers were formed with the mutant protein (**Fig. 4B**). Far UV circular dichroism measurements of TCV P38_W274A in the presence of dsRNA ligand revealed that the induced-fit binding of the RNA ligand was disabled in the mutant protein (**Fig. 4C**). While the cooperativity of the binding isotherms remained unchanged under different buffer conditions, RNA binding of TCV P38_W274A was inhibited under anionic and competitive phosphate buffer conditions (**Supplementary Fig. S6**).

The W274A mutation was shown to impair dimerization, RNA binding and subunit exchange. This suggests that the indole moiety of W274 is essential for the structural integrity of the P-domain. Furthermore, these data highlight the significance of P-domain interactions in establishing the functional TCV P38 dimer.

### Arginine residues determine the RNA-binding ability of TCV P38

A previous study described the importance of three arginine residues, R57, R74 and R84, in the TCV P38 N-terminal linker and S-domain for RNA binding (50). To assess the role of these arginine residues in the context of the structural and functional studies performed here, we recombinantly produced and purified the variants TCV P38_R57A, TCV P38_R74A and TCV P38_R84A according to established protocols (see **Fig. 1** and Materials and Methods). Far UV circular dichroism spectra of the variant proteins revealed their structural integrity, with TCV P38_R57A and TCV P38_R84A resembling closely the wildtype protein while the variant TCV P38_R74A displayed increased CD signal amplitudes in both the α-helical and β-sheet spectral regions. Interestingly, the spectrum of the variant TCV P38_R74A showed only minor changes after RNA binding. In contrast, the spectra of TCV P38_R57A and, most notably, TCV P38_R84A showed clearly measurable structural changes (**Supplementary Fig. S7**). These differences were also reflected in EMSA RNA-protein interaction studies (**Table 3**). While for TCV P38_R74A the binding affinities to siRNAs remained the same and the affinity to dsRNA_33 and to the viral UTRs (not shown) was even increased compared to the wildtype protein, this was clearly not the case with the other variants: TCV P38_R57A showed a tenfold reduction in affinity for siRNAs and microRNAs, while the K_D_ values for the dsRNA_33 and the UTRs (not shown) were unchanged compared to the wildtype protein. The variant TCV P38_R84A, which had to be purified from inclusion bodies, showed reduced affinity for both the siRNAs (∼10-fold) and dsRNA_33 (∼25 fold) (**Table 3**). No binding of this variant to longer structured RNA, such as the TCV 3’UTR, was detected (not shown).

Based on these results, we confirmed that Arg57 (in the linker between R- and S-domains) and Arg84 (in the S-domain) substantially contribute to dsRNA binding, most likely through electrostatic interactions.

### Structural studies on TCV P38

The highly pure concentrated protein was applied to crystallization experiments. RNA-free TCV P38 crystallized in the presence of 0.2 M ammonium sulfate and 22% (w/v) PEG 8000 at pH 5.7 (see Materials and Methods). The crystals diffracted to 2.9 Å in the rhombohedral space group R3, with 20 independent molecules in the asymmetric unit. Although the protein preparation consisted of dimeric RNA-free TCV P38, structure solution revealed four pentamers in a capsid-like T = 1 (60 protomers) icosahedral arrangement in the crystal (**Fig. 5A**). While all 20 S-domains could be easily located by molecular replacement, no density corresponding to the R-domains could be located and that for the P-domains was initially poor. Making use of the twenty-fold redundancy, P-domain β-strands could be identified in the averaged density in the immediate neighborhood of the S-domain shell. This in turn allowed determination of the individual P-domain dimer axis orientations, allowing local real-space averaging and completion of the model (“best” and “worst” densities are shown in **Supplementary Fig. S8**).

**Figure 5.**
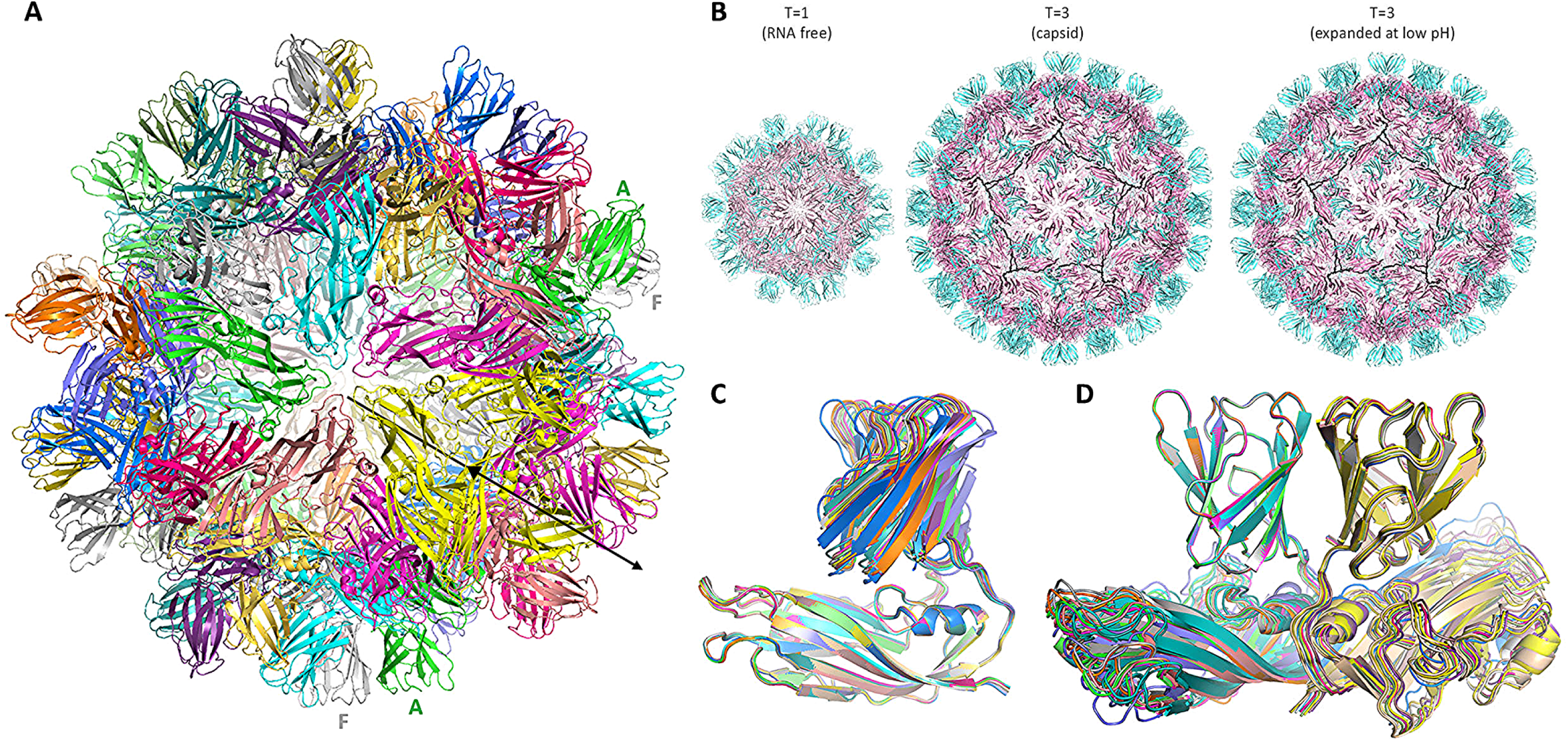
Crystal structure of RNA-free TCV P38 capsid-like icosahedron. **(A)** The protein crystallizes with twenty monomers (chains A-T, individually colored) in the asymmetric unit, which through the crystallographic threefold axis (arrow) define a 60-mer (T = 1) icosahedral capsid-like arrangement. Two copies of the best-defined P-domains (chains A, green, and F, grey) are labelled. **(B)** Comparison of the T = 1 RNA-free capsid-like structure (diameter ∼220 Å) determined here, the T = 3 virion capsid (80) (∼330 Å) and the expanded capsid observed at low pH (∼380 Å) (34) all viewed down the fivefold axis. **(C)** Superpositioning of all twenty independent copies of TCV P38 in the T = 1 icosahedron onto the S-domain of monomer A demonstrates the structural identity of the S-domains, but the P-domains adopt different orientations with respect to the S-domain. This is in contrast to superpositioning onto the P-domain of the A-monomer **(D)**, which not only shows that each P-domain has the same structure, but that the P:P-domain interaction is conserved.

The structures of the individual P38 domains superimpose well on those of the earlier published T = 3 TCV capsids [pdb code 3zxa, (35); pdb code 9qvg (80)], and with only minor adjustments of the linker segment and S-domain dimer contacts, the present structure can be aligned with the low-resolution, low pH expanded T = 3 viral capsid cryo electron microscopy reconstruction [emd1864 (34)] (**Fig. 5B**). Each TCV P38 monomer is defined in the present structure from residue D80, *i.e.* corresponding to the S-domain of A- and B-protomers in T = 3 viral capsids. The space that would be occupied by the longer C-protomer N-terminus (see **Supplementary Fig. S2**) is blocked in the T = 1 structure by adjacent protomers, indicating that N-terminal β-strand formation to construct the β-annulus structure (35) is incompatible with the present structure and is in accordance with earlier findings that N-terminally cleaved TCV P38 capsid protein forms T = 1 icosahedral particles (37,81).

Superposition of the present TCV P38 A-chain S-domain on that of the virion protomer A reveals the same pentameric arrangement in the T = 1 and T = 3 icosahedral arrangements (**Fig. 5B**). In contrast, the P-domain protrusions do not follow the S-domain T = 1 icosahedral symmetry as identified during structure solution; indeed, superposition of the TCV P38 S-domains reveals a significant variation in the relative orientation of the P-domain (**Fig. 5C**). On the other hand, superposition of the P-domains demonstrates that the P-domain dimers act as a single unit rotating about the corresponding S-domains as rigid bodies connected by the twin linker peptide segments G243-D247 (**Fig. 5D**). The P:P-dimer interface appears to be invariant between the T = 1, T = 3, and expanded forms (**Supplementary Fig. S9**).

The above comparisons revealed that the TCV capsid has the same P:P-dimer arrangement as the capsid of *Carnation Mottle Virus* (CarMV). CarMV belongs to the Alphacarmovirus genus in the *Tombusviridae* family and is only distantly related to TCV. To further evaluate this interesting observation, we submitted the complete TCV P38 sequences (UniProt P06663) and the sequences of its S-, P-, and R-domains to BLAST (63) searches in the UniProtKB (64). While the full-length TCV P38 showed sequence homologies to 34 related proteins from 10 *Tombusviridae* genera, we found 30 homologues for the S-domain, 16 for the R-domain, and remarkably, only one for the P-domain (**Table 4**; **Supplementary Table S4**). In order to perform structural comparisons, the homologous protein sequences were submitted to AlphaFold3 (66,67) and the models entered into MOE (68). These analyses revealed only minor differences in the general architecture of the respective S- and P-domains (see **Supplementary Fig. S10** and **S11**), but interestingly demonstrated the existence of two distinct clusters of *Tombusviridae* capsid proteins, referred to here as “Cluster 1” and “Cluster 2” (**Supplementary Fig. 12**; **Table 4**). Most interestingly, the AF3-based models also suggest a defined structure for the R-domain (**Fig. 6A**), supporting an earlier proposal (82) that the R-domain can adopt a relatively stable folded structure. Upon comparing the R-domain models of the viruses in Clusters 1 and 2, it became evident that there is a high degree of conservation of the primary sequence and structure within a cluster (shown for Cluster 1 in **Supplementary Fig. S13**). However, there are significant differences between the two clusters: while the R-domains of Cluster 1 viruses (to which TCV belongs) arrange into three looped α-helices, with the loop between the second and third helices containing the N-terminal GW motif (**Fig. 6A**), the R-domains of Cluster 2 consist mainly of coil structures (**Supplementary Fig. S14**). Significantly, cluster 1 contains a number of examples of Carmoviruses and Pelarspoviruses in which, analogous to TCV P38, the viral capsid protein was reported to act also as VSR, whereas viruses belonging to Cluster 2 encode separate capsid and VSR proteins as documented for Tombusviruses, Aureusviruses, and Dianthoviruses (**Table 4**; **Supplementary Table S4**; see also Discussion).

**Figure 6.**
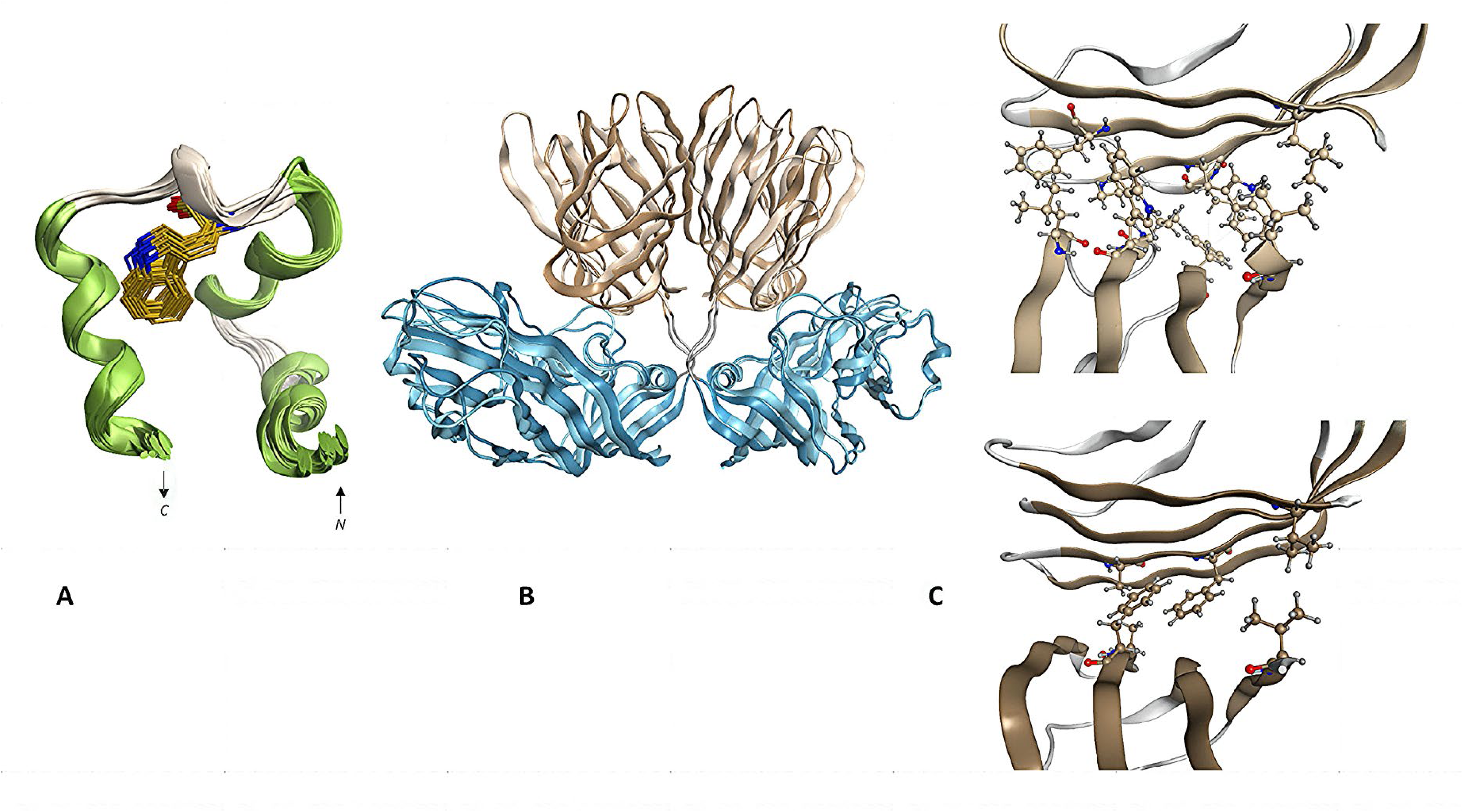
Modeling of the R-domain. Comparison of TCV P38 and TBSV P41 dimers. **(A)** Structural superposition of Cluster 1 AlphaFold3-models for the TCV P38 R-domain, with the conserved GW motif shown in stick representation. N- and C-termini are indicated. **(B)** AlphaFold3-predicted dimers of the capsid proteins from TCV (light colors) and TBSV (darker colors). The protein dimers reveal remarkable structural conservation (Cα RMSD ≈ 2.5 Å) despite substantial functional and sequence divergence (see text). **(C)** Comparison of dimer interfaces in TCV and TBSV coat proteins based on AlphaFold3-models. (*top*) The dimer interface of TCV P38 is formed by two nearly perpendicular β-sandwich structures, with extensive hydrophobic contacts between the facing sheets formed by F291, L311, V319, W321 and L349. (*bottom*) Equivalent dimer model of the TBSV P41 coat protein. Although a similar geometric arrangement is observed, fewer hydrophobic contacts of amino acid residues, A348, F350 and V377, are involved in the interaction.

**Table 4.** BLAST results for TCV P38-like capsid protein domains and reported VSRs in Tombusviridae.

| Genus (no. of sequences),<br>Cluster | BLAST search |  |  | Documented<br>VSR function |
| --- | --- | --- | --- | --- |
|  | R-domain | S-domain | P-domain |  |
| Alphacarmovirus (8), Cluster 1 | 7 | 8 | - | P37, CP (83-85) |
| Aureusvirus (4), Cluster 2 | - | 4 | - | P14 (86)** |
| Betacarmovirus (3), Cluster 1 | 3 | 3 | 1 | P38, CP (84-86) |
| Dianthovirus (2), Cluster 2 | - | 2 | - | MP (87-89) |
| Gallantivirus (1), Cluster 2 | - | 1 | - | - |
| Gammacarmovirus (2), Cluster 2 | - | 2 | - | P38, P42, CP (90-92)* |
| Macanavirus (1), Cluster 2 | - | 1 | - | - |
| Pelarspovirus (6), Cluster 1 | 5 | - | - | P37, CP (84,93) |
| Tombusvirus (5), Cluster 2 | - | 5 | - | P19 (94,95) ** |
| Tralespevirus (2), Cluster 1 | 2 | 1 | - | - |

Blast analysis of the three TCV P38 domains with UniProt reference proteomes and SwissProt. The number of sequences determined per genus is shown. Protein names, types (CP/MP – coat/capsid/movement protein), and references with reported VSR activity are indicated in the VSR column.
\* an evident VSR activity of the CP of MNSV (see also **Supplementary Table S4**) has not been determined – the published data are inconsistent
\*\* VSR function *without* capsid function

Next, we submitted the 35 sequences of the homologous *Tombusviridae* proteins to dimer modelling in AF3. Interestingly, resulting models involving solely the S- and P-domains displayed a remarkably similar architecture (**Fig. 6B**; see also **Supplementary Fig. S15**): this becomes most obvious when comparing the models of the TCV P38 (genus Betacarmovirus) with that of the P41 capsid protein of *Tomato bushy stunt virus*, TBSV (genus Tombusvirus). TBSV belongs to the Cluster 2 viruses and expresses a separate VSR, the well-characterized protein P19. Despite a very low sequence conservation between TCV P38 and TBSV P41, the FGDI-β-sheets of the P-domains of both protein dimers (**Fig. 6C**; see also **Supplementary Fig. S11**) form an extensive and structurally highly conserved interface through which the two P-domains interact at an almost right angle to each other. However, while this interface is stabilized in the TCV P38 dimer by extensive hydrophobic interactions involving at least five amino acid residues, this is not the case in the TBSV P41 dimer: here, an analogous geometric arrangement is formed by hydrophobic interactions involving only three amino acids (**Fig. 6C**). Overall, this data supports the idea that the spatial arrangement of hydrophobic amino acids within the interacting P-domains is crucial for forming and/or stabilizing the TCV P38 dimer while it functions as a VSR.

### The TCV P38 wildtype and some mutant proteins interfere with DCL and/or RISC activity

Previous studies already revealed that TCV P38 inhibits DCL and RISC activity (50) (see also Introduction). The availability of purified TCV P38 wildtype and mutant proteins enabled a deeper understanding of the underlying mechanisms of these inhibitory activities. To this end, we performed a “Dicer assay” that mimics the processing of dsRNA by DCLs *in vitro* (74,79). The assay uses a cytoplasmic extract from *Nicotiana tabacum* BY-2 cells (BYL) containing endogenous DCL4, DCL2, and DCL3 activities that generate 21-, 22-, and 24 nts siRNAs from a dsRNA substrate, respectively. Processing of a model dsRNA in the presence of TCV P38 wildtype and mutant variants (see Materials and Methods, and **Fig. 7** for details) demonstrated clear inhibition of DCL-mediated processing in the presence of the wildtype TCV P38 and the arginine variant TCV P38_R74A, but not with the other mutant proteins (**Fig. 7A**), aligning with our previous findings that TCV P38_R74A does not exhibit significant folding or dimerization defects and binds to dsRNA substrates with an affinity similar to the wildtype P38 (**Table 3**). Next, we tested the P38 variants using a “Slicer assay” that reconstitutes RISC activity *in vitro* in BYL. For this purpose, the mRNA of an AGO protein of choice is translated in the BYL in the presence of siRNA. Then the RISC reconstituted in this way is incubated with a labeled target RNA. RISC activity is measured by detecting the cleavage products of the AGO-sliced target RNA (50,74). To test for the inhibition of RISC activity, we supplemented BYL prepared in this manner with the purified TCV P38 protein variants. We observed an evident inhibition of RISC-mediated cleavage of a target RNA with the wildtype TCV P38 and the mutant protein TCV P38_R74A. Interestingly, also the mutant TCV P38_W274A protein was found to inhibit RISC catalyzed slicing. Importantly, however, inhibition was only detectable when the wildtype and mutant VSRs were introduced into the reaction prior to RISC assembly (**Fig. 7B**).

**Figure 7.**
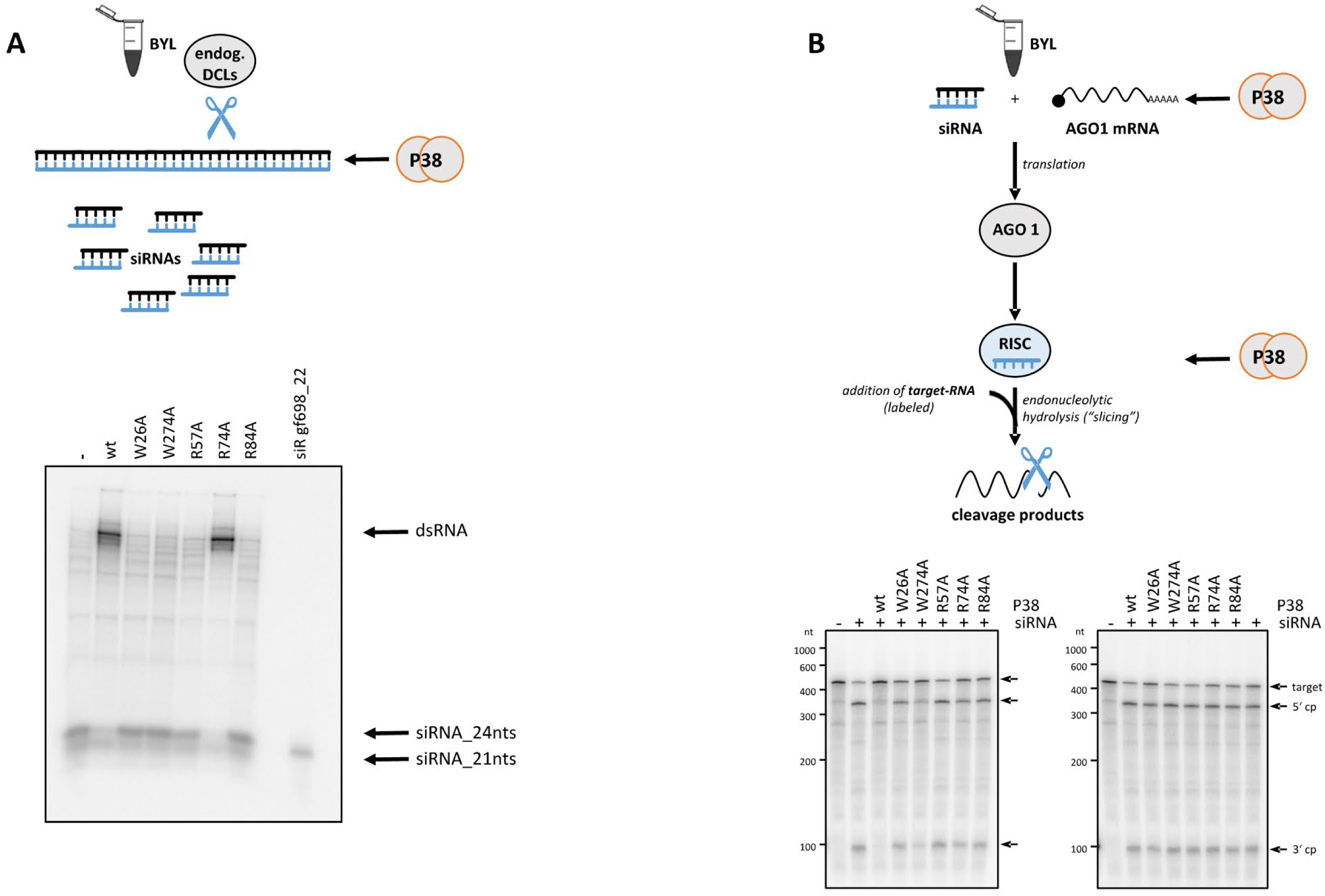
TCV P38 inhibits Dicer and RISC/Slicer activities. **(A)** Binding of dsRNA by TCV P38 and variants inhibits DCL activity by competition. (*upper panel*) Schematic representation of the “Dicer assay”. Labeled dsRNA (TCV 3’UTR) was added to BYL and processed by endogenous DCL enzymes into siRNAs. (*lower panel*) Results of “Dicer assays” performed in the presence of 100 nM (dimer) of purified TCV P38 and its variants. Processed 21 nts and 24 nts siRNAs are indicated by arrows. siR gf698_22 was used as a marker. **(B)** TCV P38 and variants affect siRNA-mediated target RNA cleavage by AGO1/RISC *in vitro*. (*upper panel*) Schematic presentation of the “Slicer assay”. *Nicotiana benthamiana* AGO1 was generated by *in vitro* translation of the respective mRNA in BYL. Translation was performed in the presence of siR gf698_21 (10 nM), thus generating RISC programmed with this siRNA. Recombinant and purified TCV P38 or respective variants (100 nM dimer) were added to the sample either at the start of the translation reaction (simultaneous incubation with the siRNA) or after RISC programming with the siRNA was complete (post-incubation). Subsequently, a ^32^P-labeled fragment of GFP mRNA containing the respective siRNA target site was added to the reaction. RNA was extracted and analyzed for RISC-mediated target cleavage by denaturing PAGE. (*lower panel left*) Results of a “Slicer assay” performed by simultaneous incubation. Target RNA and cleavage products (5’ cp, 3’ cp) are indicated by arrows. (*lower panel right*) Results of a “Slicer assay” performed by post-incubation.

In summary, these observations suggest that the inhibitory effect of TCV P38 on DCL and RISC activity is primarily mediated by the VSR competing with the cellular proteins for the RNA substrate (see Discussion). We explain the ability of TCV P38_W274A to inhibit RISC assembly by noting that, despite its altered dimerization properties, this mutant can still form functional VSR dimers with the RNA substrate (**Fig. 4**; **Table 3**).

## Discussion

Using highly purified nucleic acid-free protein fractions, we demonstrate that TCV P38 is a dimer in solution when acting as an RNA-binding protein or VSR. Analytical ultracentrifugation experiments revealed that the dimeric species is relatively stable thermodynamically, as no monomers were detectable. However, the isotope labeling experiments clearly showed that the dimer is kinetically unstable (metastable), with a very low activation barrier for the exchange reaction. These experiments also revealed that binding dsRNA prevents the exchange reaction of monomers in the dimer. In other words, interaction with dsRNA converts the protein from an exchangeable conformation to a stable, non-exchangeable one. This observation is significant because it illustrates how the VSR binds to RNA substrates. On the other hand, it sheds light on the potential transition from the VSR form to the capsid form (discussed in further detail below).

What does the TCV P38 dimer look like? Unfortunately, it was technically impossible to gain direct insight into its structure, as we were unable to obtain crystals of the dimers. Nevertheless, crystallization of the purified TCV P38 allowed us to solve the structure of a T1 capsid-like icosahedral arrangement. We assume that this was due to the high concentration of the protein preparation used in crystallization, reflecting an inherent ability of TCV P38 to self-associate to form an icosahedral arrangement even in the absence of RNA. Conversely, expression of N-terminally truncated P38 *in planta* (81) results in the formation of RNA-free T = 1 structures, whilst RNA-containing T = 3 virus-like particles are formed when the full-length protein is expressed. Thus, packaging of RNA appears to involve interplay with the N-terminal region of P38 to form the T = 3 encapsidated viral genome, presumably through an RNA-mediated ordering of the N-termini to form the β-annulus scaffold. This supports previous cryo-electron microscopy studies that allowed direct visualization of the encapsulated genomic RNA with the N-terminal region of TCV P38 as an uncoating intermediate (34).

Jellyroll structures, classically present in many viral capsid proteins, in the S- and P-domains form the scaffolding structures of the individual protein and the entire capsid, respectively (**Fig. 5**; **Supplementary Fig. S8** and **S9**). However, both data sets - derived from X-ray structural analysis of the capsid as well as AF3 modeling - highlight the P-domains as decisive determinants of TCV P38 dimerization, as noted earlier in *in planta* expression studies (81). The structural alignments of *Tombusviridae* capsid proteins and of its individual domains revealed the existence of two clusters (**Table 4**; **Supplementary Fig. S12**; **Supplementary Table S4**). Proteins of the Cluster 1 viruses function as both capsids and VSRs, with TCV P38 as prime example, whereas proteins of Cluster 2 viruses (such as TBSV P41) appear to function exclusively as capsid proteins; TBSV and other Cluster 2 viruses are reported to encode additional proteins that act as VSRs (**Table 4**). The P-domain is barely conserved at the primary sequence level, yet all *Tombusviridae* analogues appear to possess a rather similar dimer interface as suggested by the crystal structure and the modeling data comparing TCV P38 and TBSV P41 (**Fig. 6**). However, the modeling also suggests clear differences in the number of hydrophobic interactions between the dimer interfaces of the P-domains of proteins in Clusters 1 and 2. The increased number of hydrophobic contacts, such as those occurring between TCV P38 monomers, could provide additional stabilization during dimer formation outside the capsid; *i.e.*, it could provide stabilization in an environment without further capsid-specific interactions. In fact, the VSR-function of TCV P38 in solution necessitates an intrinsically stable dimer supported by an extensive hydrophobic interface. In contrast, stable interactions between TBSV coat proteins probably only occur within the structural context of the capsid.

The lack of conservation of the P-domain primary sequences suggests that it has also virus-specific functions. In the TCV capsid, for example, it could be involved in specific interactions with cellular entry receptors. Alternatively, it is possible that the P-domain interferes with RISC formation during the TCV P38 VSR activity by interacting with specific host proteins. One example of this could be the observed interactions between TCV P38 and SGS3 or AGO *via* GW273/274 (21,51,52). The mutant protein TCV P38_W274A most impressively demonstrates this, as the TCV-specific motif G273/W274 in the P-domain is disrupted, resulting in dimerization only occurring in the presence of RNA and not between mutant and wildtype monomers (**Fig. 4**). We explain the importance of G273/W274 for TCV dimerization by the fact that tryptophan is sandwiched between two lysine residues (K267 and K269) by cation-π-interactions. This idea is supported by the fact that the mutation of these residues to alanine, as modeled, leads to electrostatic repulsion and the destabilization of the P domain’s structural integrity.

The interesting flexibility of the P:P domain interaction is also evident in the crystal structure, which shows that the P:P-dimer can occupy different positions relative to the S-domains. (**Supplementary Fig. S9**). However, this cannot account for the substantial conformational changes of the VSR during RNA binding that we observed in our binding and circular dichroism studies. In fact, the data obtained with TCV P38 and the mutant proteins TCV P38_R57A and TCV P38_R84A indicate that the N-terminus of the S-domain, the hinge region between the R- and S-domains, and the R-domain are key components for the adaptive binding of dsRNA molecules of different lengths, and thus, for siRNA binding activity (see also below). Consistent with this, our modeling data suggest that the R-domain structure is only conserved among Cluster 1 *Tombusviridae* members (**Fig. 6A**; **Supplementary Figs. S12**, **S13** and **S14**). In addition to further investigating the RNA-binding determinants of the R-domain, future studies should clarify the roles of the R- and S-domains in interacting with the identified viral genome packaging signal (96).

How can one imagine the activity of TCV P38 in the course of a viral infection? After entry and uncoating, the viral genome is translated and an initial set of TCV P38 molecules is produced from subgenomic RNA. The RNA-genome is then replicated in membrane compartments, and, at the moment when the genomic RNA of the progeny is translated in the cytoplasm, the amount of TCV P38 dimers increases dramatically. High concentrations of the VSR inhibit DCL activity (50). Previously, it was unclear whether this occurred through direct interactions between TCV P38 and DCLs or through competition for RNA substrates. Unlike earlier experiments by Iki *et al.*, our complementation Dicer assays (**Fig. 7A**) revealed that only the wildtype protein and the TCV P38_R74A variant inhibit DCL processing. Both proteins bind RNA with an extremely high affinity (**Table 3**), clearly supporting the hypothesis that TCV P38 primarily inhibits DCL activity by competing for RNA and in keeping with the dissociation constant for TCV P38 – dsRNA binding in the pico- to nanomolar range (**Table 3**; summarized in **Fig. 8**).

**Figure 8.**
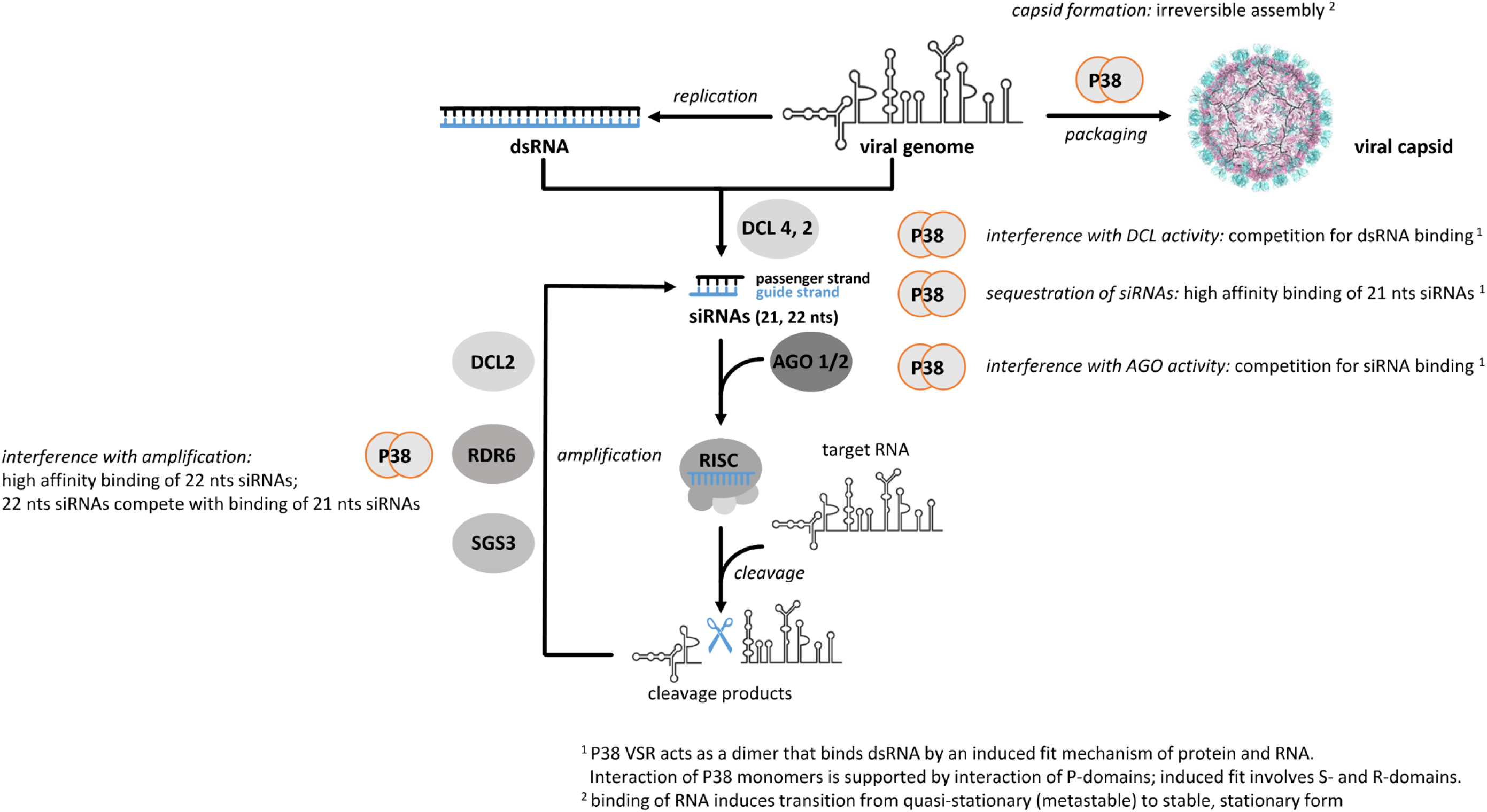
Scheme of the antiviral RNA silencing process and formation of the viral capsid. Dicer-like proteins (DCLs), in plant mainly DCL4 and DCL2, process double-stranded dsRNA elements of a target RNA (viral genome or mRNA) into small interfering siRNAs, of which mainly 21 nts and 22 nts siRNAs are involved in antiviral RNAi (see text). Argonaute endonucleases (AGO) incorporate one siRNA strand, *in planta*, mainly AGO1 and AGO2 are antivirally active. *Via* base-pairing, this guides AGO and other proteins as an RNA induced silencing complex (RISC) to the target RNA. The target RNA is inactivated by cleavage or translation inhibition (not shown). siRNAs are amplified *via* the activity of RNA-dependent RNA polymerases and DCLs (indicated here for RDR6, in cooperation with SGS3 and DCL2). Mechanisms of TCV P38 protein activities characterized during this study are indicated.

Despite this affinity for TCV P38, it can be assumed that certain quantities of 21 nts siRNAs are still produced in the infected cells. These are also bound by TCV P38, preventing the formation of active RISC (**Fig. 2**). Using again the established *in vitro* system, this time to perform Slicer assays in the presence of wild-type and mutant VSRs, we found that in addition to the wildtype protein also the variants TCV p38_R74A and TCV p38_W274A inhibit RISC-mediated cleavage of a target RNA. Importantly, this was only observed when the VSRs were introduced before RISC assembly occurred. Therefore, we conclude that the siRNAs bound to RISC are inaccessible to the viral proteins. Thus, in line with an earlier report by Iki *et al.* (50), the inhibitory effect of TCV P38 on RISC activity appears to be primarily due to competition with the RISC-enclosed AGO protein for siRNA binding. Interestingly, also TCV P38_W274A inhibits RISC assembly. Although this variant has a significantly higher dissociation constant with the RNA substrate and exhibits a significantly reduced ability to form functional VSR dimers compared to the wildtype protein, the latter ability appears to be crucial for successful competition with AGO.

Our competition experiments revealed that TCV P38 binds 21 nts siRNAs with high affinity, yet these siRNAs can still be displaced by RNAs of a higher molecular weight (**Fig. 2D**). This can be explained simply by the fact that the high molecular weight RNAs offer more binding sites as polymer substrate for the protein. This is of great importance given that 21 nts siRNAs are generated during initial phases of the infection, while 22 nts siRNAs are predominantly produced in a second phase of the infection (51). We postulate that replacement of initial 21 nts siRNAs associated with TCV P38 molecules with newly generated 22 nts siRNAs maintains P38 as a constant and effective active VSR throughout the plant’s RNA silencing response (**Fig. 8**).

TCV P38 showed the highest affinity for higher molecular weight dsRNAs (**Table 1**). This finding is consistent with previous observations that the RNA size is an important determinant of the TCV assembly process (96). Evidently, in the late phase of infection when viral protein and genome concentrations are high, TCV P38 dimers are more likely to bind to viral RNAs. As shown here, this process occurs cooperatively, forming further intermediates with increasing occupancy, which ultimately leads to capsid formation. The packaging signal is located in the TCV P38 protein’s own coding region (96); it may act as a feedback control that blocks further translation, for example (96). Accordingly, our data suggest the following sequence of events during capsid formation. Given our observations of the VSR dimer transitioning from a quasi-stationary to a stationary form during RNA binding, we can assume this sequence is irreversible. When the metastable TCV P38 dimers associate with the RNA, the binding energy exceeds a threshold value at a critical oligomer concentration or “seeding”. The reverse reaction (disassembly – endothermic) does not occur due to additional intra- and intermolecular interactions during the formation of the regular structure (assembly – exothermic). As a result, RNA-containing capsids form (**Fig. 8**).

## Supporting information

Supplemental data complete

## Acknowledgements

We thank Christine Hamann and Katja Rostowski for technical assistance.

## Author contributions

Dennis Arendt: Methodology, Investigation, Validation, Formal Analysis Selvaraj Tamilarasan: Methodology, Investigation, Validation, Formal Analysis

Ralph Peter Golbik: Methodology, Conceptualization, Investigation, Validation, Formal Analysis, Data Curation, Writing – Original Draft, Supervision

Iris Thondorf: Methodology, Conceptualization, Investigation, Software, Validation, Formal Analysis, Data Curation

Julian Bender: Methodology, Investigation, Validation, Formal Analysis, Data curation, Funding Aquisition

Hauke Lilie: Methodology, Investigation, Validation

Torsten Gursinsky: Methodology, Conceptualization, Investigation, Validation

Christoph Parthier: Methodology, Investigation, Validation, Formal Analysis, Data curation Carla Schmidt: Conceptualization, Validation, Resources, Data Curation, Supervision, Funding Acquisition

Milton T. Stubbs: Conceptualization, Validation, Resources, Data Curation, Visualization, Writing – Review & Editing

Sven-Erik Behrens: Conceptualization, Validation, Resources, Data Curation, Writing – Original Draft, Writing – Review & Editing, Supervision, Project Administration, Funding Acquisition

## Supplementary data

Supplementary Data are available online.

## Data availability

The data underlying this article are available in the article and in its online supplementary material

