## Supplemental data complete for "A flexible Janus head: molecular determinants of a viral protein’s RNAi suppressor and capsid forming activities"

**Supplementary data**

**Supplementary Materials and Methods**

**Recombinant production and purification of $^{14}N$- and $^{15}N$-TCV P38 and variants.** The fusion protein $His_6$-SUMO-TCV P38 and the variants were recombinantly produced in *Escherichia coli* BL21-CodonPlus (DE)-RIPL. Bacteria were grown in kanamycin-containing LB medium at 37 °C until an optical density of about 1.0. Gene expression was induced by addition of 1 mM isopropyl-1-thio-β-D-galactopyranoside (IPTG, Roth) and proceeded at 25 °C overnight. The production of the $^{15}N$-isotopically labelled protein was performed by using the same expression system, but bacteria were grown in kanamycin-containing M9 medium (minimal medium) supplemented by $^{15}NH_4Cl$ (Sigma-Aldrich) as nitrogen source. Harvested bacteria were re-suspended in 250 mL lysis 50 mM potassium phosphate, 400 mM NaCl, 100 mM KCl, 10 mM imidazole, 10 % glycerol, 0.1 % Triton X-114, pH 7.4 containing protease inhibitors (cOmplete™, Roche). Lysis was performed by addition of lysozyme (1.5 mg per g bacteria, Sigma-Aldrich), incubation on ice for 20 min and afterwards for 20 min at room temperature. Cell rupture was completed by dispersing using an Ultra Turrax and treatment in a French Press (Gaulin). Nucleic acids were digested by addition of 10 µg $mL^{-1}$ DNase I (Roche) and 3 mM $MgCl_2$ to the suspension and stirring at room temperature for 30 min. Ultracentrifugation (L8-60M, BECKMAN) at 30,000 rpm for 60 min at 10 °C yielded a clear supernatant and a pellet of cell debris. The supernatant was thoroughly dialysed against 50 mM sodium phosphate, 500 mM NaCl, 1 mM DTT, pH 7.6 and applied to a first affinity chromatography (HisTrap™ FF, 5 mL, GE Healthcare). After washing the gradient elution of the fusion protein was performed with 50 mM sodium phosphate, 500 mM NaCl, 1 mM DTT, pH 7.6 containing 500 mM imidazole. Fractions containing the fusion protein were again dialysed against 50 mM sodium phosphate, 500 mM NaCl, 1 mM DTT, pH 7.6 for removal of imidazole. Cleavage of the fusion protein was done by incubation of 200 µL $His_6$-SUMO-protease ULP1 (305 µM) with 60 mL solution at room temperature for 30 min. Afterwards, the solution was applied to a second affinity chromatography (HisTrap™ FF, 5 mL, GE Healthcare). Following elution, the target protein was dialysed against 50 mM TRIS, 100 mM NaCl, 1 mM TCEP, pH 7.6. Precipitated material was removed by ultracentrifugation (L8-60M, BECKMAN) at 35,000 rpm for 60 min at 10 °C. The supernatant was concentrated to about 10 mL using concentrators (MWCO 20 kDa for monomers or 50 kDa for dimers, Sartorius) and applied to gel filtration (Superdex 75 16/60, GE Healthcare) for elution with 50 mM TRIS, 100 mM NaCl, 1 mM TCEP, pH 7.6. Fractions with target protein were then applied to affinity / cation exchange chromatography (HiTrap™ Heparin HP, 5 mL, Cytiva). The gradient elution was performed with 50 mM TRIS, 1 M NaCl, 1 mM TCEP, pH 7.6. This affinity / cation exchange chromatography was essential to separate the dimer fractions containing truncated protein from correctly assembled native dimers. The preparation of $^{15}$N-isotopically labelled TCV P38 was carried out according to this established protocol as well (**Fig. 1A**). All recombinant proteins were in soluble form except the variant TCV P38_R84A. Here, the inclusion bodies were obtained after ultracentrifugation (L8-60M, BECKMAN) at 30,000 rpm for 30 min at 10 °C and treated / washed according to the protocol established by RUDOLPH and LILIE (1,2). Afterwards the protein was denatured by solubilisation in 100 mM TRIS, 6 M GdmHCl, 100 mM DTT, 1 mM EDTA, pH 8.0 at 25 °C. Precipitation of contaminating nucleic acids was conducted by dropwise adding concentrated HCl to pH 2.0 at 4 °C. Removal of the precipitant was performed by ultracentrifugation (L8-60M, BECKMAN) at 35,000 rpm for 60 min at 8 °C. The supernatant was dialysed against 20 mM TRIS, 40 mM NaCl, 6 M urea, pH 7.6 and supplied to anion exchange chromatography (HiPrepQ 16/10, GE Healthcare). Protein elution succeeded in a linear gradient using buffer 8 (20 mM TRIS, 940 mM NaCl, 6 M urea, pH 7.6). For cleavage of $His_6$-SUMO-TCV P38_R84A by $His_6$-SUMO-protease ULP1 refolding was initiated by dialysis against 50 mM sodium phosphate, 500 mM NaCl, 1 mM DTT, pH 7.6 for at least 15 hours. After cleavage of the refolded protein by $His_6$-SUMO-protease ULP1 (mentioned above) the solution was applied to affinity chromatography (HisTrap™ FF, 5 mL, GE Healthcare). The material was again denatured in 20 mM TRIS, 40 mM NaCl, 6 M urea, pH 7.6 due to precipitation, refolded in 50 mM TRIS, 1.5 M NaCl containing 0.5 M L-arginine, pH 7.6 and applied to gel filtration (Superdex 75 16/60, GE Healthcare) for elution with 50 mM TRIS, 100 mM NaCl, 1 mM TCEP, pH 7.6. Protein containing fractions were pooled and separated by affinity / cation exchange chromatography on Heparin as described above. Target proteins were generally analysed by SDS-PAGE and mass spectrometry (**Fig. 1B**). The protein concentrations were determined by absorbance measurements to ensure removal of nucleic acids and aggregate detection. The extinction coefficient of TCV P38 (molecular weight 38139.06 Da) and the arginine variants (molecular weight 38053.95 Da) was 57,410 $M^{-1}$ $cm^{-1}$, that of the tryptophan variants TCV P38_W26A and TCV P38_W274A (molecular weight 38023.92 Da) was 51,910 $M^{-1}$ $cm^{-1}$ and of TCV P38_W26A_W274A (molecular weight 37908.79 Da) was 46,410 $M^{-1}$ $cm^{-1}$.

**Origin, generation and modification of RNAs.** The siR gf698_22 used in this study was earlier described (3). The siR gf698_21 and siR gf698_24 siRNAs were derived from the entire original sequence (*GFP*). The sequences of the miRNAs from *Arabidopsis thaliana* (*At*) and *Nicotiana benthamiana* (*Nb*) were derived as described in PERTERMANN *et al.* (4). The sequences of TCV 3'UTR were derived from the *Turnip crinkle virus* genome deposited in the database. RNA oligonucleotides (**Supplementary Table S3**) to generate the here-examined RNAs were purchased from Biomers (Ulm, Germany). Since phosphorylation was earlier shown to increase the binding affinity of RNAs by VSRs (5) all oligonucleotides were 5'-

phosphorylated using T4 polynucleotide kinase (Thermo Fisher Scientific) and standard procedures. For radiolabeling, 25 µCi of [$\gamma$-$^{32}$P] ATP (3,000 Ci mmol$^{-1}$) was added to the phosphorylation reaction. For sRNA annealing, the phosphorylation reactions were stopped by adding EDTA and reactions of two complementary oligonucleotides combined. After heating at 94 °C for 3 min, the mixture was cooled at a rate of 1 degree per min to 25 °C. Hybridized RNA-duplexes were then purified using illustra™ microSpin™ G-25 columns (GE Healthcare) as suggested by the manufacturer. To generate the radiolabeled TCV 3'UTR as target RNA for the DCL assay, the cDNA of the 253 nts fragment was amplified by PCR with the T7 promotor sequence included in the (respective) forward primer. Transcription of the (respective) target RNAs was performed from the PCR products by T7 RNA polymerase in the presence of 0.5 µCi µL$^{-1}$ [$\alpha$-$^{32}$P] CTP (3,000 Ci mmol$^{-1}$) under standard conditions.

**Circular dichroism studies of TCV P38 and variants.** Chirooptical properties of proteins and nucleic acids are distinctive in far and near ultraviolet spectrum. To detect conformational changes during complex formation of TCV P38 and variants with RNAs spectra of the single components and of the complexes were recorded. Spectra of complexes were overlaid with the theoretical spectra resulting from the summation of the respective single component spectra. This approach allowed an illustration of conformational changes of both parts during complex formation. Circular dichroism spectra were recorded on a dichrograph J-810 from JASCO using cuvettes with 1 mm optical path length. Measurements were carried out at 20 °C in 50 mM TRIS, 100 mM NaCl, 1 mM TCEP, pH 7.6 at protein concentrations typically of 2-3 µM dimer (4-6 µM monomer) and at siRNA concentrations adjusted in equimolar manner. Spectra were accumulated 40 times. For data investigation and spectra comparison molar ellipticity values were plotted considering the monomer concentration of the proteins.

**Supplementary Tables**

**Supplementary Table S1.**

**DNA oligonucleotides**

| primer name | sequence | purpose |
|---|---|---|
| pSUMOp38f1 | GACCATGAAGACCGTGGTATGGAAAATGATCCTAGAGTCC | forward PCR primer for *TCV* P38 cloning into vector pET-SUMO-adapt |
| pSUMOp38r1 | CCGCTCGAGTTAAATTCTGAGTGCTTGCCATTTACC | reverse PCR primer for *TCV* P38 cloning in vector pET-SUMO-adapt |
| p38GW1mut2f | GTGGCAGAAGAAGGGCGCGTCAACCCTAACCAGC | forward PCR primer for site-directed mutagenesis of *TCV* P38 (W26A) |
| p38GW1mut2r | GCTGGTTAGGGTTGACGCGCCCTTCTTCTGCCAC | reverse PCR primer for site-directed mutagenesis of *TCV* P38 (W26A) |
| p38GW2mut1f | GGACAGCTGGGGCGGAGCACGATTGTC | forward PCR primer for site-directed mutagenesis of *TCV* P38 (W274A) |
| p38GW2mut1r | GACAATCGTGCTCCGCCCCAGCTGTCC | reverse PCR primer for site-directed mutagenesis of *TCV* P38 (W274A) |
| p38R57mut1f | CCTGTGCAGAAAGTGACTGCACTGAGTGCTCCGG | forward PCR primer for site-directed mutagenesis of *TCV* P38 (R57A) |
| p38R57mut1r | CCGGAGCACTCAGTGCAGTCACTTTCTGCACAGG | reverse PCR primer for site-directed mutagenesis of *TCV* P38 (R57A) |
| p38R74mut1f | GTTACCACCCAGCCTGCGGTCTCTACTGCCAG | forward PCR primer for site-directed mutagenesis of *TCV* P38 (R74A) |
| p38R74mut1r | CTGGCAGTAGAGACCGCAGGCTGGGTGGTAAC | reverse PCR primer for site-directed mutagenesis of *TCV* P38 (R74A) |
| p38R84mut1f | CAGGGACGGCATAACCGCAAGCGGTTCTGAACTG | forward PCR primer for site-directed mutagenesis of *TCV* P38 (R84A) |
| p38R84mut1r | CAGTTCAGAACCGCTTGCGGTTATGCCGTCCCTG | reverse PCR primer for site-directed mutagenesis of *TCV* P38 (R84A) |
| T7TCVoa3f | CCCTAATACGACTCACTATATGACATTGTTCTACGAGAAGG | forward PCR primer for synthesis of *TCV* 3'UTR |
| TraTCVoa3r | CTGAGTGCTTGCCATTTACC | reverse PCR primer for synthesis of *TCV* 3'UTR |

The plasmid pTCV containing the cDNA of the entire genome of *Turnip crinkle virus* was a gift from Dr. Anne Simon (University of Maryland, USA).

**RNA oligonucleotides**

| name | sequence | purpose |
|---|---|---|
| siR gf698 21U gs | uaguucauccaugccaugugu | guide strand of gf698_21 nts siRNA |
| siR gf698 21U ps | acauggcauggaugaacuaua | passenger strand of gf698_21 nts siRNA |
| siR gf698 22U gs | uaguucauccaugccaugugua | guide strand of gf698_22 nts siRNA |
| siR gf698 22U ps | cacauggcauggaugaacuaua | passenger strand of gf698_22 nts siRNA |
| siR gf698 24U gs | uaguucauccaugccauguguaau | guide strand of gf698_24 nts siRNA |
| siR gf698 24U ps | uacacauggcauggaugaacuaua | passenger strand of gf698_24 nts siRNA |
| syn_dsRNA_33 nts fs | cuaagaugcucgcugccaaugaacgacccucccuaa | forward strand of syn_dsRNA_33 nts |
| syn_dsRNA_33 nts rs | uuagggagggucguucauuggcagcgagcaucu | reverse strand of syn_dsRNA_33 nts |
| *At*miRNA_162 gs | ucgauaaaccucugcauccag | guide strand of *At*miR162 (isoforms a and b) |
| *At*miRNA_162 ps | ggaggcagcgguucaucgauc | passenger strand of *At*miR162 (isoforms a and b) |
| *At*miRNA_168 gs | ucgcuuggugcaggucgggaa | guide strand of *At*miR168 (isoforms a and b) |
| *At*miRNA_168 ps | cccgccuugcaucaacugaau | passenger strand of *At*miR168 (isoform a) |
| *At*miRNA_403a gs | uuagauucacgcacaaacucg | guide strand of *At*miR403 and *Nb*miR403 (both isoforms) |
| *At*miRNA_403a ps | uguuuugugcuugaaucuaauu | passenger strand of *At*miR403 |
| *Nb*miR403a | uguuugugcgugaaucugaca | passenger strand of *Nb*miR403 (isoform a) |
| *Nb*miR403b | uguuugugcgugauucugaca | passenger strand of *Nb*miR403 (isoform b) |

**Supplementary Table S2. Statistics of structure refinement of TCV P38**

| **Data collection** | |
|---|---|
| X-ray source<br>wavelength (Å)<br>space group | BESSY BL14.2<br>0.9184<br>R3 |
| cell parameter<br>a,b,c (Å)<br>α,β,γ (°) | <br>204.18, 204.18, 470.81<br>90.00, 90.00, 120.00 |
| resolution (Å) | 30.0-2.9<br>(3.1-2.9) |
| completeness (%) | 99.9 (90.5) |
| total reflections<br>unique reflections | 896078<br>162094 (26071) |
| multiplicity | 5.5 (5.2) |
| $R_{merge}$ [1] | 19.3 (167.8) |
| $I/\sigma(I)$ | 9.7 (1.0) |
| $CC_{1/2}$ | 99.0 (35.1) |
| Wilson B-factor | 51.4 |
| structure refinement<br>molecules per asymmetric unit | <br>20 |
| R values (%)<br>$R_{work}$<br>$R_{free}$ | <br>17.4<br>19.3 |
| twinning fraction [2] | 0.62 |
| number of atoms | 41357 |
| average B factors ($Å^2$)<br>all atoms<br>S domain atoms<br>P domain atoms | <br>73.6<br>56.9<br>98.5 |
| Rmsd<br>bond lengths (Å)<br>bond angles (°) | <br>0.003<br>0.65 |
| Ramachandran (%)<br>favored<br>allowed<br>outlier | <br>96.44<br>3.52<br>0.04 |
| Molprobity clashscore | 6.49 |
| PDB accession code | 9RP8 |

Values for highest resolution shell are given in parentheses.
[1] $R_{merge} = \sum_{hkl}\sum_i |I_i(hkl) - \langle I(hkl)\rangle| / \sum_{hkl}\sum_i I_i(hkl) \times 100$
[2] merohedral twinning operator: h, -h-k, -l

**Supplementary Table S3. Mass spectrometry analysis of TCV P38 and TCV P38_W27A**

| protein | siRNA (siR gf698) concentration | protein concentration | Figure |
|---|---|---|---|
| TCV P38 | | 4.7 μM $^{14}$N-TCV P38 | 1C, 3A |
| TCV P38 | | 5.9 μM $^{15}$N-TCV P38 | 3A |
| TCV P38 | <1 μM 21 nts[1] | 2.1 μM $^{14}$N-TCV P38 | 1C, 3B |
| TCV P38 | <1 μM 21 nts[1] | 2.2 μM $^{15}$N-TCV P38 | 3B |
| TCV P38 | <1 μM 22 nts[1] | 4.5 μM $^{14}$N-TCV P38 | 1C, 3C |
| TCV P38 | <1 μM 22 nts[1] | 5.7 μM $^{15}$N-TCV P38 | 3C |
| TCV P38 | | 2.6 μM $^{14}$N-TCV P38, 2.7 μM $^{15}$N-TCV P38 | 3A |
| TCV P38 | <1 μM 21 nts[1] | 1.2 μM $^{14}$N-TCV P38, 1.2 μM $^{15}$N-TCV P38 | 3B |
| TCV P38 | <1 μM 22 nts[1] | 2.6 μM $^{14}$N-TCV P38, 2.7 μM $^{15}$N-TCV P38 | 3C |
| TCV P38_W274A | | 22.8 μM TCV P38_W274A | 4B |
| TCV P38_W274A | 1.45 μM 21 nts | 20.9 μM TCV P38_W274A | 4B |
| TCV P38 and TCV P38_W274A | | 17 μM TCV P38_W274A, 1.2 μM $^{15}$N-TCV P38 | 4B |
| TCV P38 and TCV P38_W274A | 1.6 μM 21 nts[1] | 17 μM TCV P38_W274A, 1.2 μM $^{15}$N-TCV P38 | 4B |

[1] Free RNA was removed by filtration before analysis.

**Supplementary Table S4. BLAST results and structural features of plant viral coat proteins inferred from TCV P38 domains and AlphaFold3 models**

| | BLAST search | | | Domain boundaries | | | AlphaFold3 pTM (ipTM) [a] | | Documented VSR function |
|---|---|---|---|---|---|---|---|---|---|
| **Genus** | **R-domain** | **S-domain** | **P-domain** | **R-domain** | **S-domain** | **P-domain** | **monomer** | **dimer** | |
| **Alphacarmovirus (ACV)** | | | | | | | | | |
| *Adonis mosaic virus* (AdMV) | ✓ | ✓ | | 11-47 | 80-242 | 248-346 | 0.73 | 0.75 (0.69) | |
| *Angelonia flower break virus* (AnFBV) | ✓ | ✓ | | 5-41 | 80-243 | 249-350 | 0.74 | 0.90 (0.89) | |
| *Carnation mottle virus* (CarMV) | | ✓ | | 12-48 | 81-243 | 249-347 | 0.70 | 0.66 (0.63) | P37, CP (6) binds siRNAs, prevents RISC incorporation, no impairment of siRNA production |
| *Calibrachoa mottle virus* (CbMV) | ✓ | ✓ | | 8-43 | 77-239 | 245-341 | 0.71 | 0.75 (0.69) | |
| *Honeysuckle ringspot virus* (HnRSV) | ✓ | ✓ | | 8-43 | 77-240 | 246-343 | 0.72 | 0.71 (0.67) | |
| *Nootka lupine vein clearing virus* (NLVCV) | ✓ | ✓ | | 11-46 | 78-240 | 246-344 | 0.73 | 0.82 (0.79) | |

| | | | | | | | | |
|---|---|---|---|---|---|---|---|---|
| *Pelargonium flower break virus* (PFBV) | ✓ | ✓ | 8-43 | 77-240 | 246-344 | 0.75 | 0.80 (0.77) | P37, CP (7,8) binds siRNAs, prevents RISC incorporation, no impairment of siRNA production |
| *Saguaro cactus virus* (SgCV) | ✓ | ✓ | 9-44 | 78-241 | 247-343 | 0.75 | 0.77 (0.73) | |
| **Aureusvirus (AurV)** | | | | | | | | |
| *Cucumber leaf spot virus* (CLSV) | | ✓ | | 92-260 | 266-377 | 0.76 | 0.76 (0.74) | |
| *Elderberry aureusvirus 1* (ElAV1) | | ✓ | | 95-260 | 266-367 | 0.76 | 0.73 (0.71) | |
| *Johnsongrass chlorotic stripe mosaic virus* (JCSMV) | | ✓ | | 74-240 | 246-355 | 0.79 | 0.81 (0.76) | |
| *Pothos latent virus* (PoLV) | | ✓ | | 91-255 | 261-362 | 0.75 | 0.64 (0.62) | P14 (9) binds long dsRNA and siRNA duplexes; independently of capsid function |
| **Betacarmovirus (BCV)** | | | | | | | | |

| | | | | | | | | | |
|---|---|---|---|---|---|---|---|---|---|
| *Cardamine chlorotic fleck virus* (CCFV) | ✓ | ✓ | ✓ | 7-42 | 82-244 | 250-350 | 0.73 | 0.66 (0.52) | |
| *Hibiscus chlorotic ringspot virus* (HCRSV) | ✓ | ✓ | | 7-42 | 82-245 | 251-344 | 0.76 | 0.73 (0.69) | P38, CP (10) suppresses initiation step of RNA silencing, presumably by interfering with the RDR6/SGS3 system |
| *Japanese iris necrotic ring virus* (JINRV) | ✓ | ✓ | | 7-43 | 83-244 | 250-351 | 0.76 | 0.78 (0.76) | |
| *Turnip crinkle virus* (TCV) | | | | 5-40 | 80-242 | 248-350 | 0.74 | 0.64 (0.60) | P38, CP (8,11) binds dsRNA & siRNA, inhibits siRNA production and RISC loading |
| **Dianthovirus (DiaV)** | | | | | | | | | |
| *Carnation ringspot virus* (CRSV) | | ✓ | | | 47-213 | 219-332 | 0.76 | 0.45 (0.25) | |
| *Red clover necrotic mosaic virus* (RCNMV) | | ✓ | | | 46-211 | 217-329 | 0.78 | 0.86 (0.82) | MP(12-14) assumed interaction with host factors |
| **Gallantivirus (GalV)** | | | | | | | | | |

| | | | | | | | | |
|---|---|---|---|---|---|---|---|---|
| *Galinsoga mosaic virus* (GaMV) | | ✓ | | 56-221 | 227-330 | 0.77 | 0.73 (0.71) | |
| **Gammacarmovirus (GCV)** | | | | | | | | |
| *Melon necrotic spot virus* (MNSV) | | ✓ | | 95-260 | 266-381 | 0.76 | 0.81 (0.77) | P38, P42, CP (15-17)[b]<br>RNA sequestration? |
| *Pea stem necrosis virus* (PSNV) | | ✓ | | 49-215 | 221-332 | 0.80 | 0.81 (0.77) | |
| **Macanavirus (MCV)** | | | | | | | | |
| *Furcraea necrotic streak virus* (FNSV) | | ✓ | | 53-221 | 227-336 | 0.84 | 0.83 (0.79) | |
| **Pelarspovirus (PelV)** | | | | | | | | |
| *Clematis chlorotic mottle virus* (ClCMV) | ✓ | | 7-41 | 79-234 | 240-336 | 0.72 | 0.55 (0.40) | |
| *Jasmine mosaic-associated virus 2* (JMaV) | | | 7-41 | 79-234 | 240-338 | 0.72 | 0.87 (0.86) | |
| *Pelargonium chlorotic ring pattern virus* (PCRPV) | ✓ | | 7-41 | 79-234 | 240-336 | 0.72 | 0.81 (0.72) | |
| *Pelargonium line pattern virus* (PLPV) | ✓ | | 7-41 | 78-233 | 239-337 | 0.73 | 0.86 (0.82) | P37, CP (8,18)<br>siRNA sequestration |

| | | | | | | | | |
|---|---|---|---|---|---|---|---|---|
| *Pelargonium ringspot virus* (PelRSV) | ✓ | | 6-40 | 78-233 | 239-337 | 0.73 | 0.89 (0.86) | |
| *Rosa rugosa leaf distortion virus* (RrLDV) | ✓ | | 7-41 | 79-234 | 240-338 | 0.74 | 0.81 (0.76) | |
| **Tombusvirus (TBV)** | | | | | | | | |
| *Artichoke mottled crinkle virus* (AMCV) | | ✓ | | 102-267 | 273-378 | 0.75 | 0.74 (0.71) | |
| *Cucumber necrosis virus* (CNV) | | ✓ | | 92-258 | 264-370 | 0.76 | 0.88 (0.84) | |
| *Cymbidium ringspot virus* (CymRSV) | | ✓ | | 98-263 | 269-370 | 0.75 | 0.76 (0.74) | |
| *Lisianthus necrosis virus* (LNV) | | ✓ | | 102-267 | 273-378 | 0.75 | 0.72 (0.70) | |
| *Tomato bushy stunt virus* (TBSV) | | ✓ | | 102-267 | 273-378 | 0.75 | 0.77 (0.74) | P19 (5,19) siRNA sequestration, independently of capsid function |
| **Tralespevirus (TSV)** | | | | | | | | |
| *Gompholobium virus A* (GomVA) | ✓ | ✓ | 7-43 | 79-243 | 249-346 | 0.74 | 0.56 (0.48) | |
| *Trailing lespedeza virus 1* (TLV1) | ✓ | | 9-45 | 81-244 | 250-345 | 0.73 | 0.63 (0.49) | |

BLAST searches were performed with the individual domain sequences of the TCV P38 coat protein in UniProt Reference Proteomes and Swiss-Prot databases. From left to right: the first three columns indicate the presence of hits for the respective domains (✓), grouped by genus. Note that these hits were obtained only when the individual domain sequences were used as queries. In contrast, all proteins listed in the table were retrieved when we used the full-length TCV P38 sequence indicating overall sequence similarity across genera. The subsequent three columns list the domain boundaries as determined by structural and sequence alignments of the corresponding AlphaFold3 (AF3) models using the Molecular Operating Environment (MOE). For the R-domain, only the AF3-models listed here display a clear topology of three $\alpha$-helices; the remaining, predicted structures are largely disordered in this region. The final two columns show the predicted TM (pTM) and interface predicted by template modeling (ipTM) scores of the AF3 monomer and dimer models. All pTM values are in the reliable range (> 70 %) but slightly reduced due to the presence of a long linker between the R- and S-domains and/or intrinsic disorder in the N-terminal region. AlphaFold3 consistently predicted the same dimer topology and relative orientation of the monomers across all runs (**Fig. S13**). The interface predicted TM-scores (ipTM) mostly fell into the intermediate confidence range (0.6–0.8), with some values >0.8 and one <0.6. While ipTM scores in the 0.6–0.8 range indicate increased uncertainty in the precise inter-monomer orientation, the reproducibility of the overall topology across models supports the plausibility of the predicted dimer arrangement.

[a] ipTM values are calculated only for the dimer

[b] Reports on the RNA silencing suppressor (VSR) activity of the MNSV coat protein (CP) are inconsistent. While classical assays did not reveal any suppression activity (14), later studies described RNA-binding properties and RSS-like effects under specific experimental conditions such as overexpression or organelle targeting (15,16). If MNSV CP does display any suppressor activity, it appears to be weak and/or conditional, and does not resemble the strong CP-based VSRs characterized in other Tombusviruses, such as TCV P38 or PLPV P37.

### Supplementary Figures and Legends

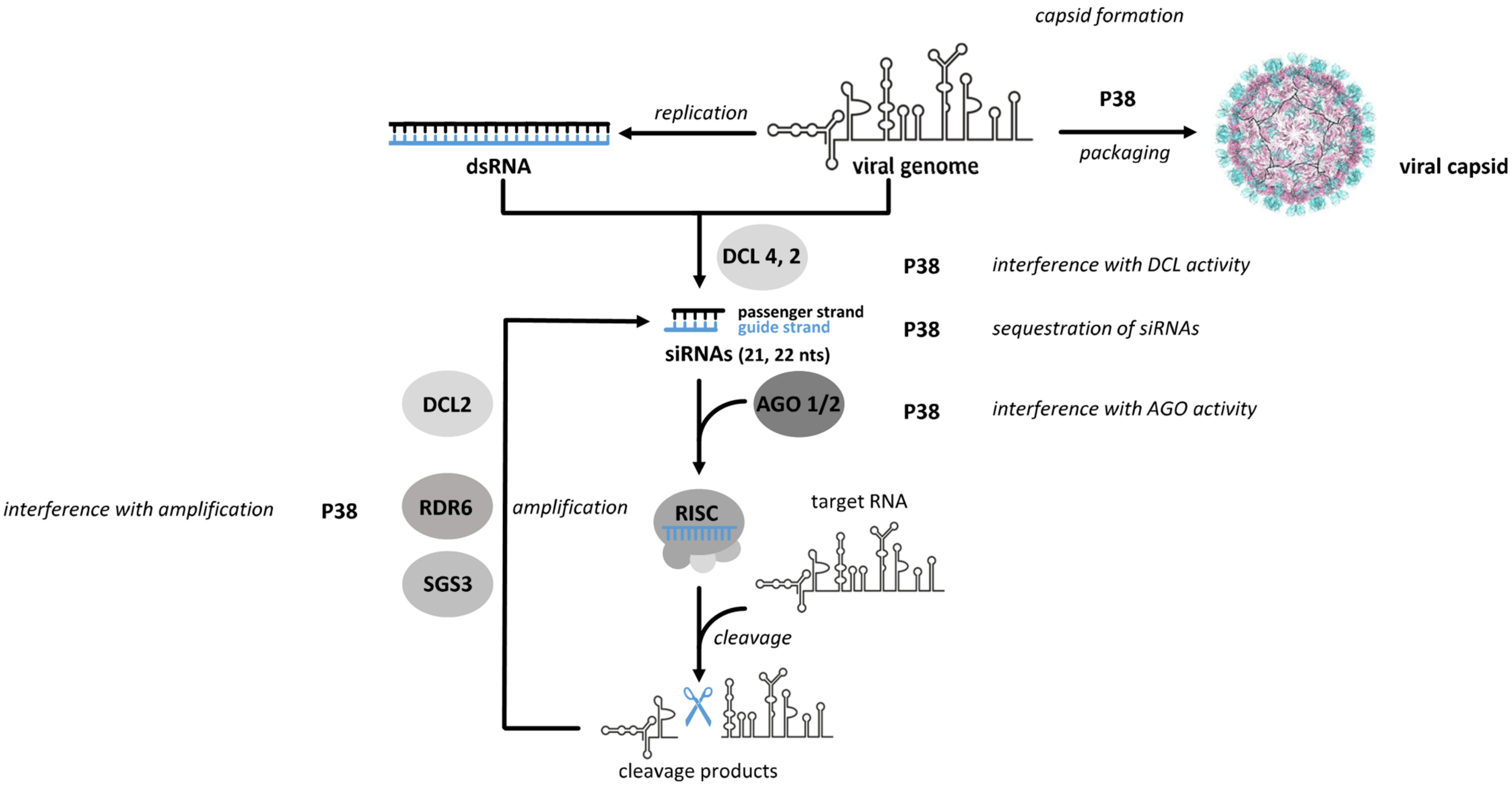

**Supplementary Figure S1. Scheme of the antiviral RNA silencing process and formation of the viral capsid.** Dicer-like proteins (DCLs), in plant mainly DCL4 and DCL2, process double-stranded RNA elements of a target RNA (viral genome or mRNA) into small interfering siRNAs. Involved in antiviral RNAi are mainly 21 nts and 22 nts siRNAs (see text). Argonaute endonucleases (AGO) incorporate one siRNA strand. Antivirally active in plant are mainly AGO1 and AGO2. *Via* base-pairing, this guides AGO and other proteins as a RNA induced silencing complex (RISC) to the target RNA. The target RNA is inactivated by cleavage or translation inhibition (not shown). siRNAs are amplified *via* the activity of RNA-dependent RNA polymerases and DCLs (indicated here for RDR6, in cooperation with SGS3 and DCL2). The activities of the TCV P38 protein that were known prior to this study are indicated.

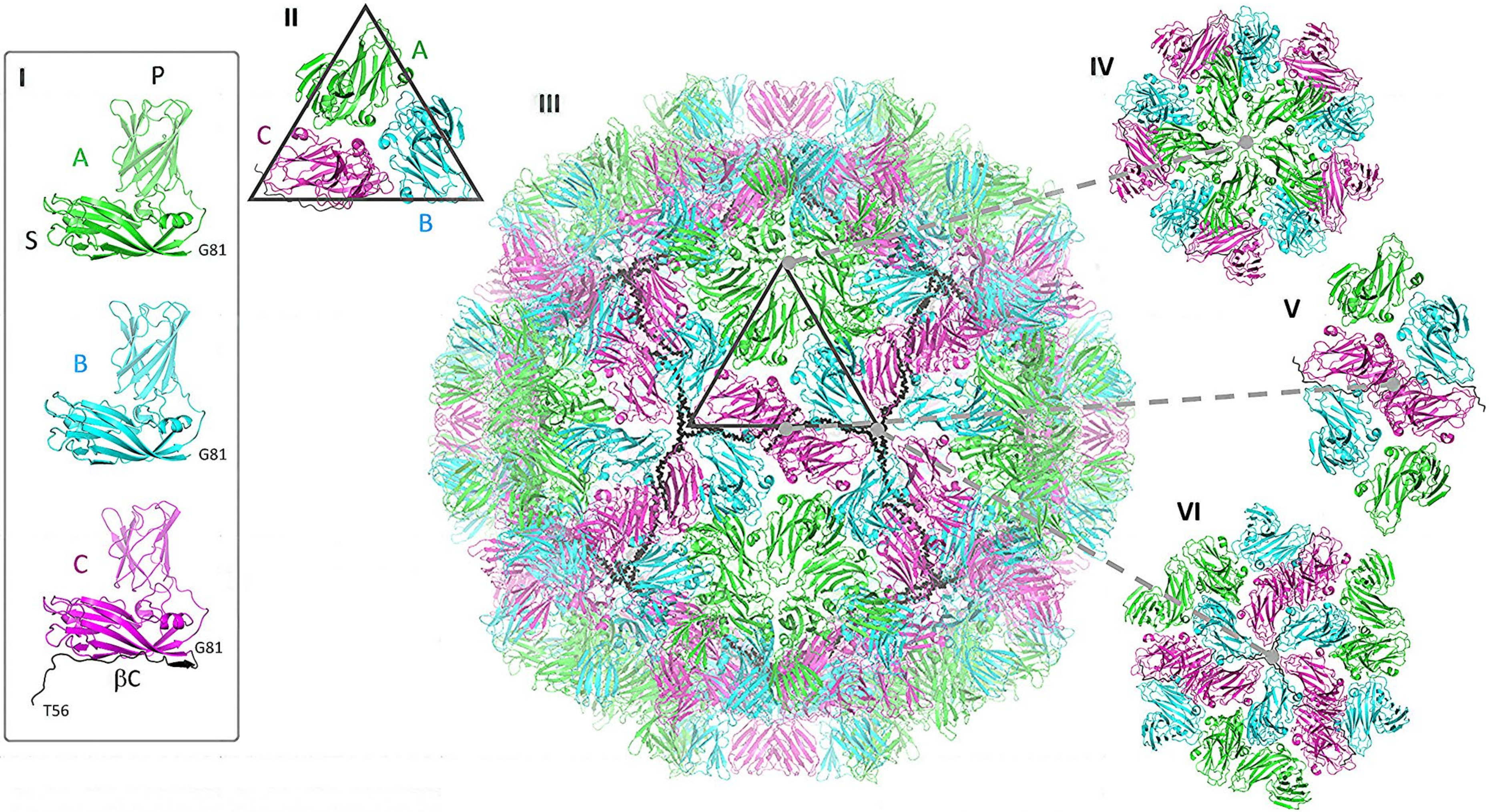
I
P
A
S
G81
B
G81
C
G81
βC
T56
II
A
C
B
III
IV
V
VI

**Supplementary Figure S2. Structure of TCV P38 in viral capsids. (I)** In mature TCV virus capsids (20-22), the TCV P38 protein forms a protein coat embedding the viral RNA. The TCV P38 protein can be divided into three domains – an internal N-terminal R-domain not seen in the capsids (as this does not follow the icosahedral symmetry of the coat) and two jelly roll domains S- (which form the shell) and P- (which protrude from the outer surface). The protein is found in three quasi-equivalent environments A (green), B (cyan) and C (magenta) in a triangular arrangement **(II)** that make up the faces of a regular (T = 3) icosahedron **(III)**. The A-conformers arrange as a pentamer around the fivefold axis **(IV)**, whereas the C-conformers are found at the twofold axis **(V)**. The N-termini of the C-conformers are extended in a b-strand-like conformation (black, see also **IC**) and form a β-annulus around the threefold axis **(VI)**. The B- and C- conformers arrange around the latter to present a pseudo sixfold axis. In the capsid **(III)**, the β-annuli make up a dodecahedral scaffold (black) that interlock the individual pentamers.

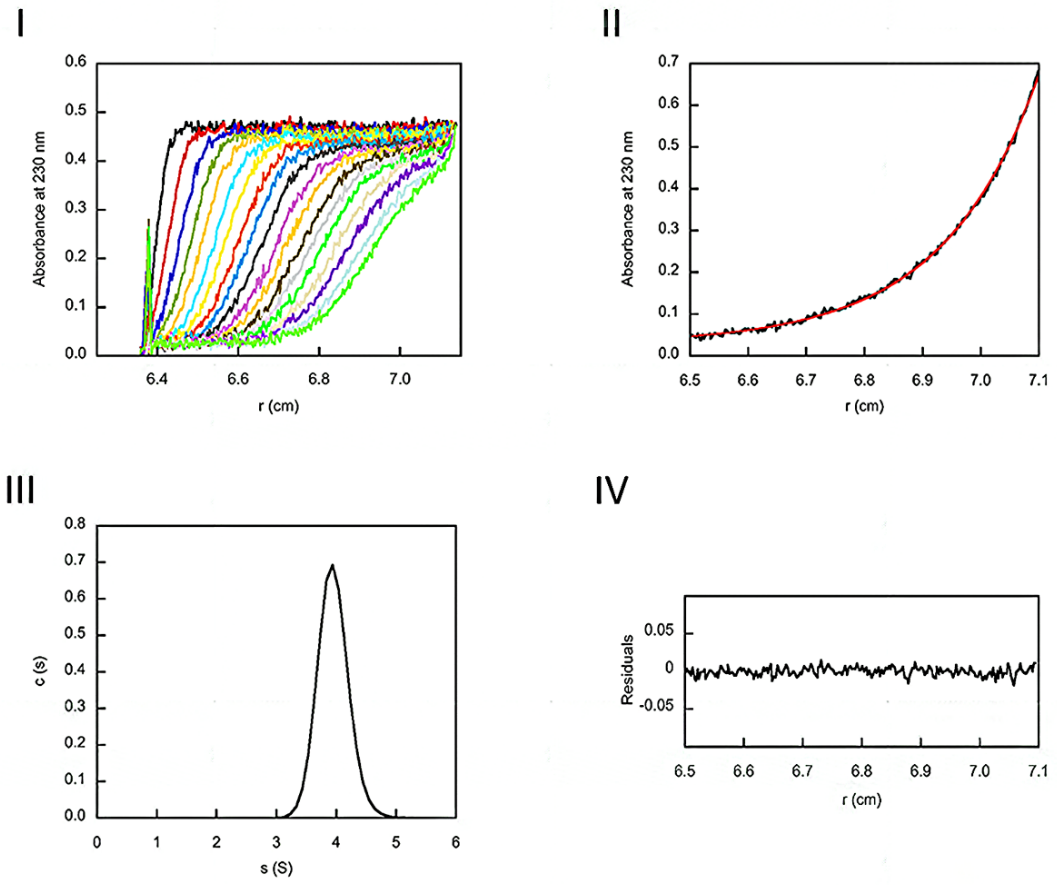


**Supplementary Figure S3. Analytical ultracentrifugation of TCV P38. (I)** Sedimentation velocity of the protein monitored at 230 nm every 10 min during 3 h. **(II)** Fitting (red line) of the sedimentation equilibrium (black line) revealed a molecular mass $M_r = 70.3$ kDa that corresponded to a dimer. **(III)** Data analysis using the software Sedfit resulted in a homogenous species with a sedimentation coefficient s = 3.9 S. **(IV)** Deviation of the fit to the data.

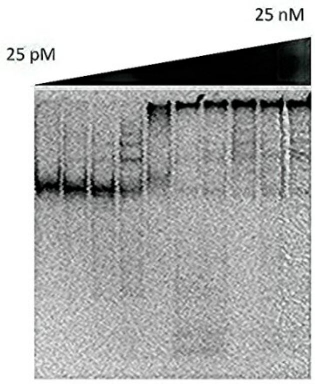


**Supplementary Figure S4. Binding of TCV P38 to TCV 3'UTR.** Analysis of TCV P38 binding to TCV 3'UTR by EMSA revealed a stepwise process and the formation of several complexes of bound RNA in the analytical gel. Binding isotherms were fitted with a cooperativity value of 2 indicating multiple independent binding events. The protein concentration indicated on top of the gel refers to the dimer.

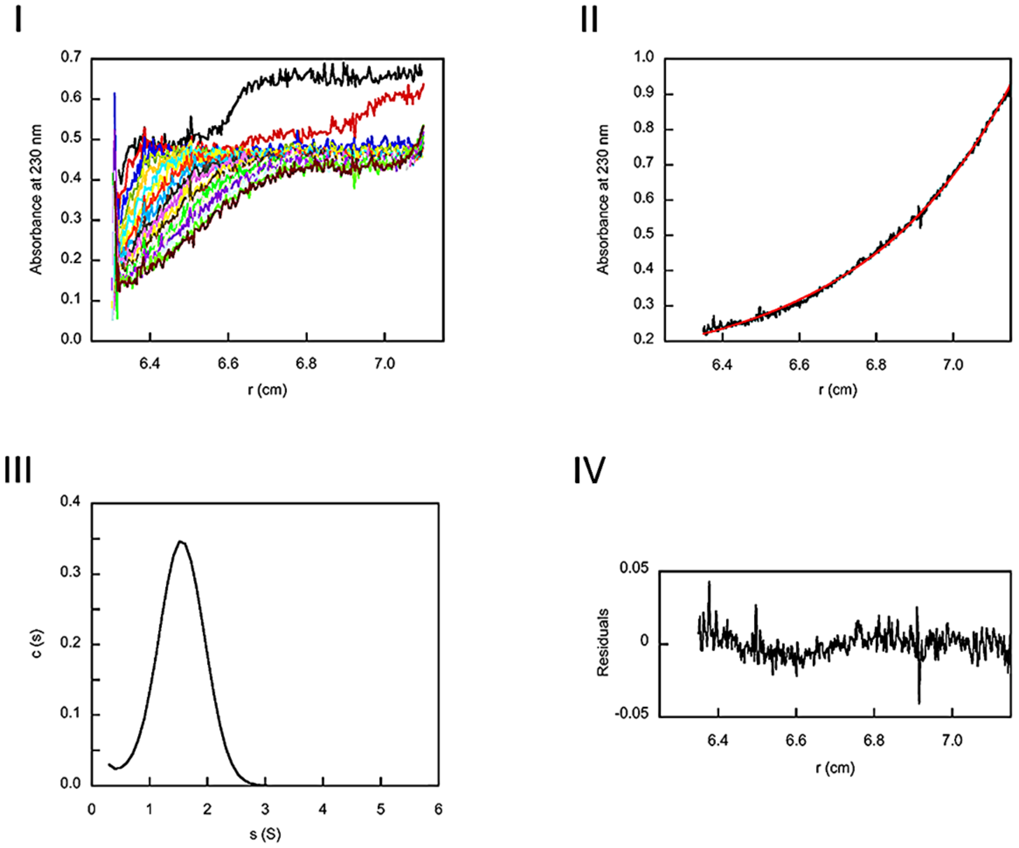


**Supplementary Figure S5. Analytical ultracentrifugation of TCV P38_W274A. (I)** Sedimentation velocity of the protein monitored at 230 nm every 10 min during 3 h. **(II)** Fitting (red line) of the sedimentation equilibrium (black line) revealed a molecular mass $M_r$ = 30 kDa that corresponded to a monomer. **(III)** Data analysis using the software Sedfit resulted in a homogenous species with a sedimentation coefficient s = 1.55 S. **(IV)** Deviation of the fit to the data.

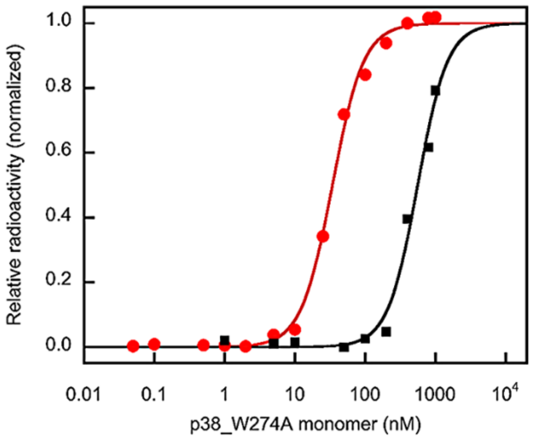


**Supplementary Figure S6. Binding of TCV P38_W274A to dsRNA is strongly affected by the ionic environment.** Binding of TCV P38_W274A to siR gf698_21 in cationic and anionic buffer was monitored by EMSA, respectively. The binding isotherms were measured in 50 mM TRIS, 0.1 M NaCl, 1 mM TCEP, pH 7.6 (red circle) or 50 mM sodium phosphate, 0.1 M NaCl, 1 mM TCEP, pH 7.6 (black square) and fitted according to equation 2 with a cooperativity factor of 2. The transitions displayed the same slope. $K_D$ values are summarized in **Table 3** and reveal an about tenfold impairment of binding dsRNA by the variant in phosphate buffer due to competition.

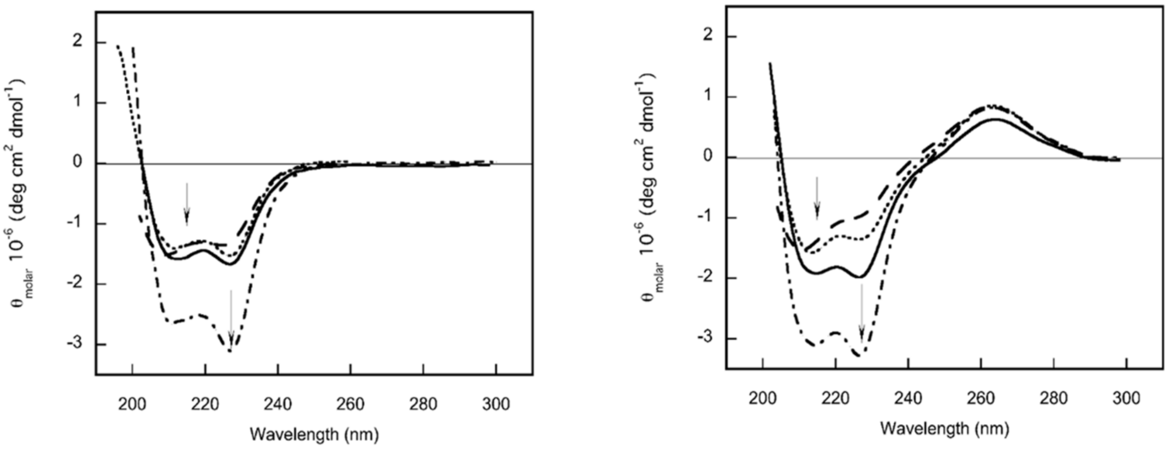


**Supplementary Figure S7. RNA binding by arginine variants of TCV P38.** (*left*) The far and near UV circular dichroism spectra of the variants displayed the same shape as the wildtype spectrum (extrema marked by arrows) with α-helical secondary structure elements and a negative extremum at 227 nm (TCV P38 – black solid line, TCV P38_R57A – black dotted line, TCV P38_R74A – black dot-dashed line, TCV P38_R84A – black dashed line). The spectrum of TCV P38_R74A displayed a double signal amplitude and a more pronounced ellipticity at 227 nm in comparison to 210 nm. (*right*) The binding of siR gf698_21 to the arginine variants in equimolar manner (considering the protein concentration as dimer) led to changes in their secondary structures. For TCV P38 (black solid line) and TCV P38_R74A (black dot-dashed line) the overall helicity increased with the biggest signal amplitude at 227 nm. In contrast, the variants TCV P38_R57A (black dotted line) and TCV P38_R84A (black dashed line) revealed some structural perturbations with a smaller signal amplitude at 227 nm in comparison to 210 nm. The residues R57 and R84 thus were determined to be critical for binding dsRNA (see also **Table 3**).

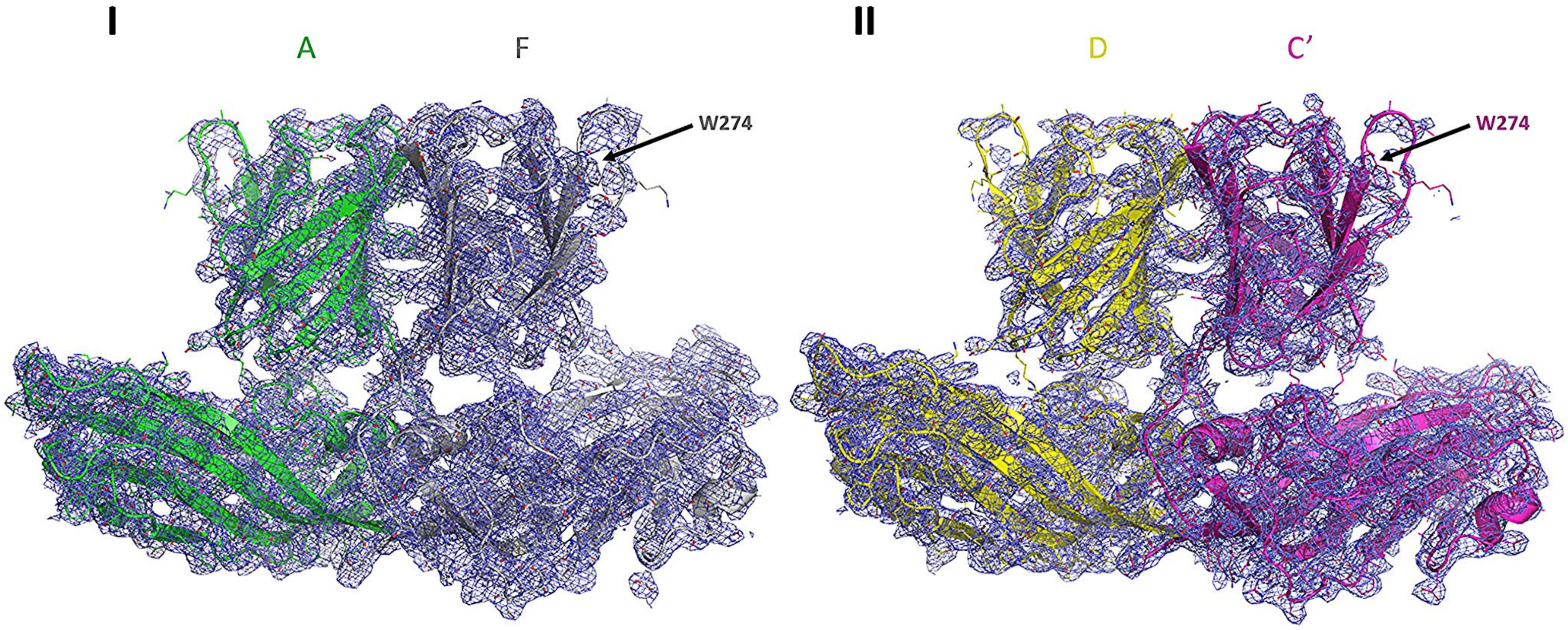
I
A
F
W274
II
D
C′
W274

**Supplementary Figure S8. Experimental electron density distributions (2Fo-Fc, contoured at 1s) for RNA-free TCV P38 dimers. (I)** Density for the S-domains is well defined and continuous in all monomers of the asymmetric unit, and that for the P-domain is most complete for the dimer AF. **(II)** Although the dimer positioning is unambiguous for all monomers, the outer loops in *e.g.* dimer DC' (crystal symmetry related to C) are less well defined, particularly in the neighborhood of $G^{273}W^{274}$ (indicated).

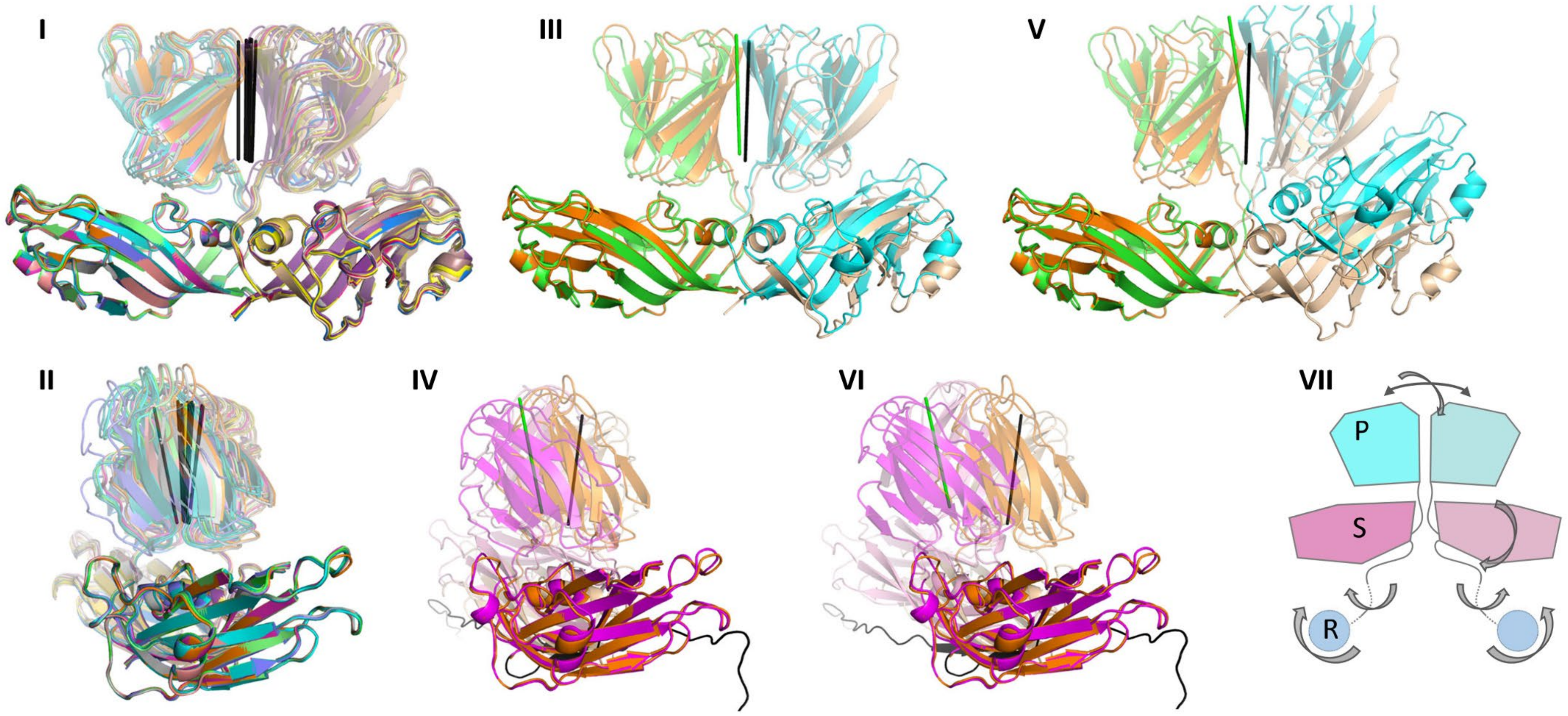
I
III
V
II
IV
VI
VII
P
S
R

**Supplementary Figure S9. The RNA-free T = 1 icosahedral crystal structure reveals that P:P-domain dimers act as a single unit, whereas S:P- and S:S-domain arrangements exhibit a degree of freedom. (I)** Superposition of T = 1 dimers from the present structure on the A monomer S-domain indicates that S:S-contacts are identical as would be expected in a T = 1 icosahedron. The P:P-domains, which superimpose as dimers (**Fig. 5D**), adopt different positions with respect to the S-domain(s); P:P-domain dimer axes displayed as black rods. **(II)** As **(I)**, rotated 90° about the vertical S:S-domain twofold axis. P:P-dimers GO and HT display the most extreme positions. **(III)** AB-dimer from the viral capsid (blue/cyan; pdb code 9qvf) superimposed on the H-monomer S-domain (orange) from the present RNA-free structure with corresponding T-monomer; orientation as in **(I)**. Note the tilt of the P:P-dimer axis (AB green, HT black) and repositioning of the B/T S-domains. **(IV)** Comparison of the viral CC'-dimers (magenta/pink) with the present HT-dimer, orientation as in **(II)**. **(V)** Comparison of the HT-dimer with the AB-dimer from the low pH expanded virus (pdb code 3zx9) (21) and **(VI)** with the CC'-dimer from the same reconstruction. **(VII)** Schematic representation demonstrating the degrees of freedom available within TCV P38 dimers, using one S-domain as reference. R-domains are not visible in any TCV P38 structures investigated to date as these do not follow the icosahedral symmetries of the coat proteins in the crystals / capsids.

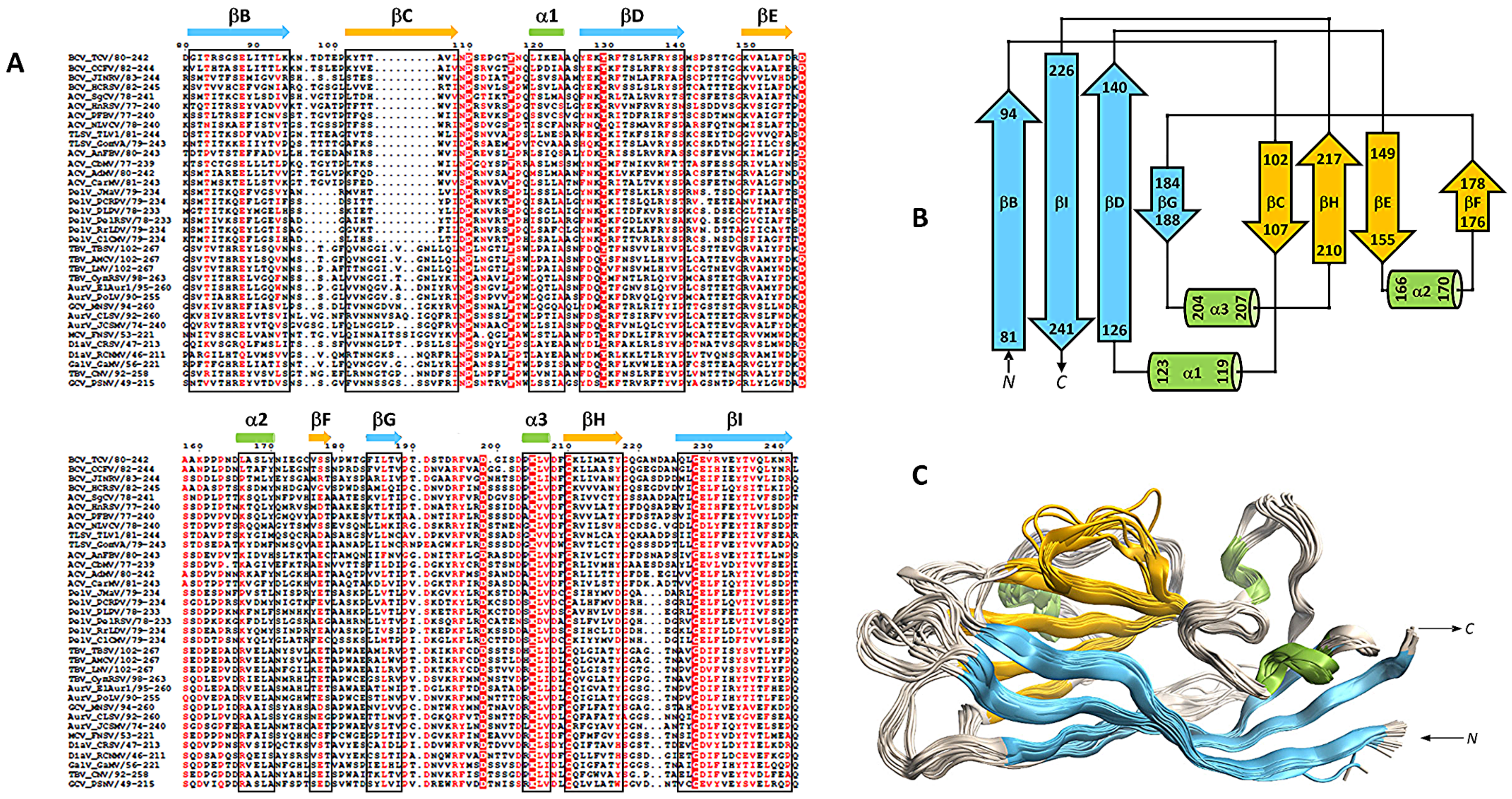
A
B
C
βB
βC
α1
βD
βE
α2
βF
βG
α3
βH
βI
N
C

**Supplementary Figure S10. Structural and sequence alignment of the S-domains of 35 homologous coat proteins. (A)** Multiple sequence alignment of the S-domains from 35 homologous plant viral coat proteins (see **Table 4**), generated in MOE (Molecular Operating Environment) based on combined sequence and structural alignment. The alignment was visualized with ESPript using BLOSUM62 as similarity matrix and a global similarity score of 0.2. Fully conserved residues are shown in white on red, strongly conserved residues in bold red, and semi-conserved residues in plain red. Secondary structure elements are indicated above the alignment as arrows (β-strands) and cylinders (α-helices), and the corresponding region in the alignment - based on the TCV P38 structure - is highlighted with black boxes. **(B)** Topology diagram of the S-domain from the AlphaFold3 (AF3) model of TCV P38. **(C)** Structural superposition of the S-domains from all 35 AF3 models, as generated in MOE. The color coding for α-helices and β-strands is consistent across all panels. Sequence and structural comparisons indicate that the S-domains of *Tombusviridae* coat proteins segregate into two major clusters (see text and **Supplementary Fig. S12**). While conserved residues are maintained across both groups - likely reflecting their importance for the integrity of the capsid fold - proteins of the second cluster (Tombus-, Aureus-, Gammacarmo-, Diantho-, Gallanti- and Macanaviruses) exhibit a distinct insert in β-strand C.

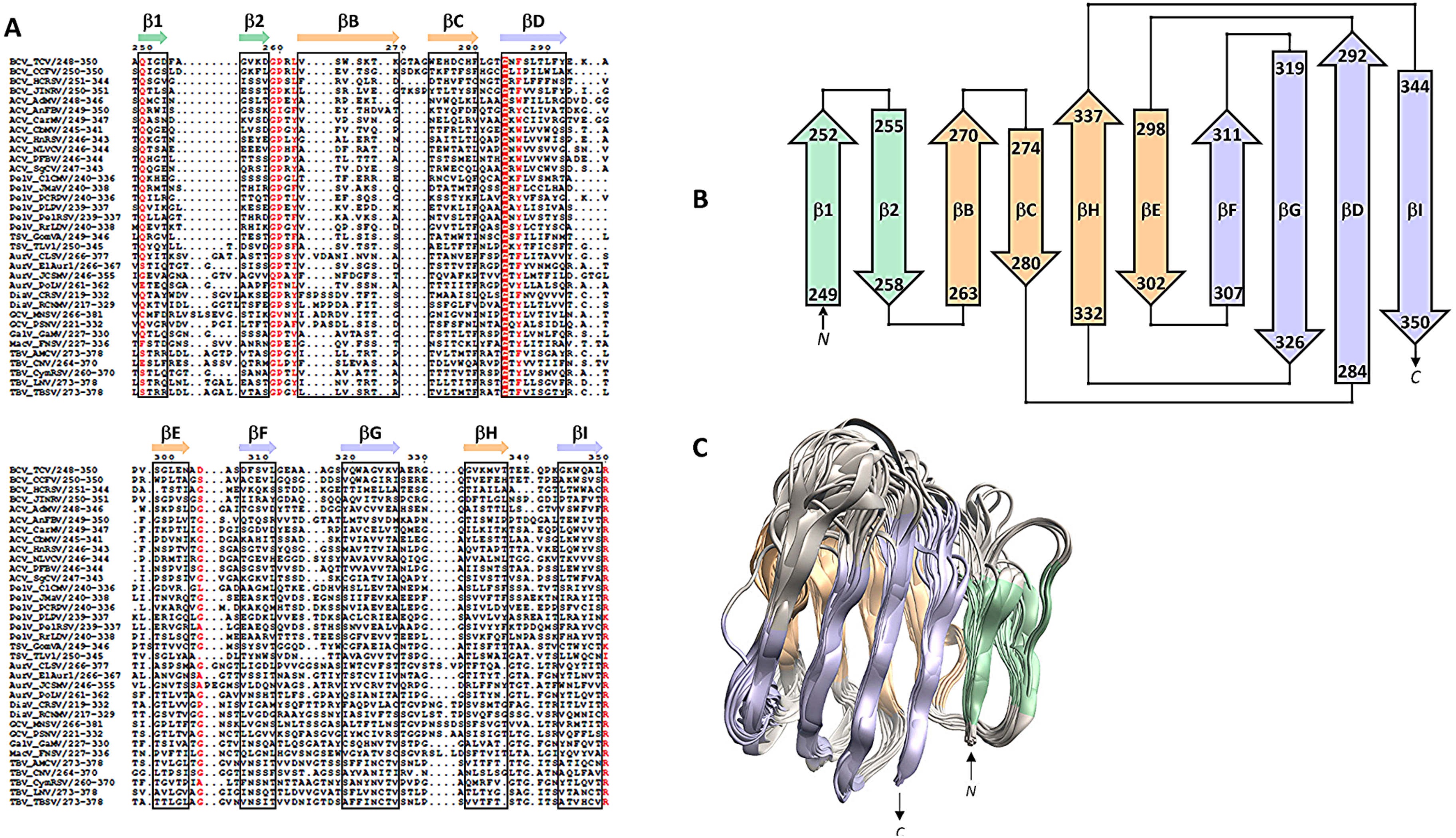
A
β1
β2
βB
βC
βD
βE
βF
βG
βH
βI
B
252
249
255
258
270
263
274
280
337
332
298
302
311
307
319
326
292
284
344
350
N
C
C
N
C

**Supplementary Figure S11. Structural and sequence alignment of the P-domains of 35 homologous coat proteins.** **(A)** Multiple alignment of the P-domains from the same 35 homologous coat proteins shown in **Supplementary Fig. S10**, generated using the same procedure (MOE-based sequence and structure alignment, ESPript visualization with BLOSUM62 matrix and global similarity score of 0.2). Conserved residues and secondary structure elements are displayed as in **Supplementary Fig. S10**. Few conserved residues are observed. **(B)** Topology diagram of the P-domain from the AlphaFold3 (AF3) model of TCV P38. **(C)** Structural superposition of all 35 AF3 models shows that, despite their low sequence similarity, the P-domains adopt similar overall folds. Structural differences are most pronounced in the distal part of the domain (upper region in panel C), whereas the region adjacent to the S-domain (lower part in panel C) is more conserved in structure. The colouring of β-strands is consistent across all panels.

**S-domain**

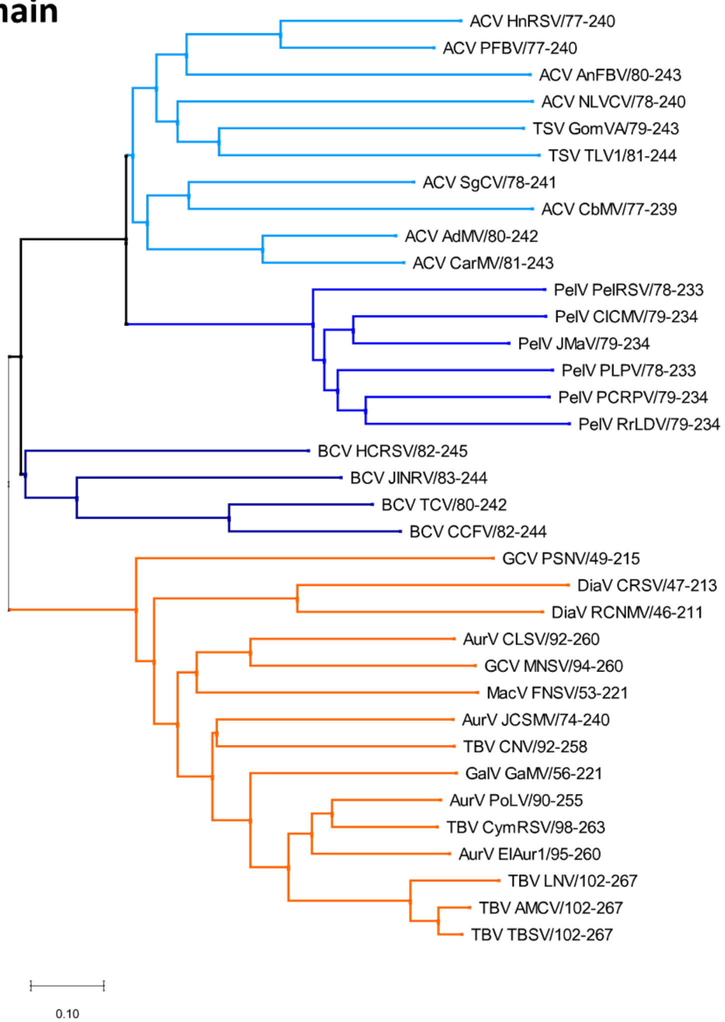


**P-domain**

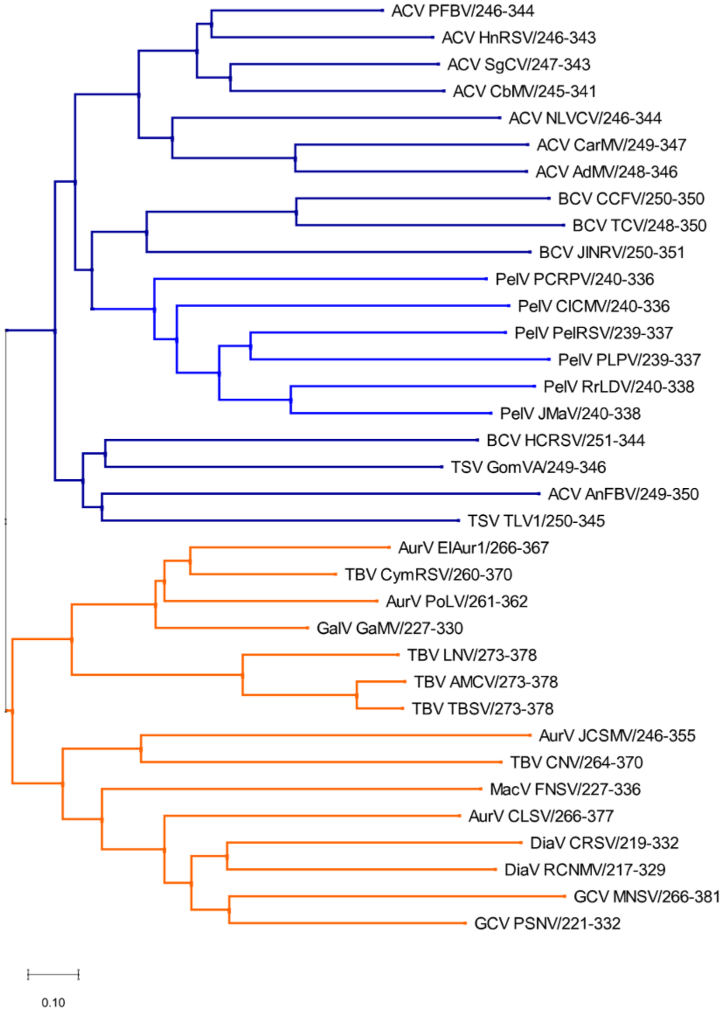

**Supplementary Figure S12. Structure- and sequence-based alignment of *Tombusviridae* capsid proteins using AlphaFold3-predicted models reveals two clusters of viruses.** Phylogenetic analysis was performed using the neighbor joining algorithm included in the MEGA program, version 11.0.13 (23), based on a multiple alignment incorporating both structural and sequence information. The AlphaFold3-predicted models of the 35 homologous coat proteins (see **Table 4**) were structurally superimposed and aligned using MOE (Molecular Operating Environment). Structural alignment was performed *via* minimization of Cα root mean square deviation (RMSD), followed by global sequence alignment using the BLOSUM62 substitution matrix. The phylogenetic trees reflect structural similarity and conservation, and are not strictly based on classical sequence-based evolutionary inference. To minimize structural uncertainties in the alignment, only models with a mean pTM greater than 70 were used. The tree topology remained stable across 10,000 bootstrap replicates. Because the focus was on qualitative structural similarity, bootstrap values and distances were not included in the diagram. The clades of Cluster 1 are colored in different shades of blue according to the genus, and the clades of Cluster 2 are colored in orange.

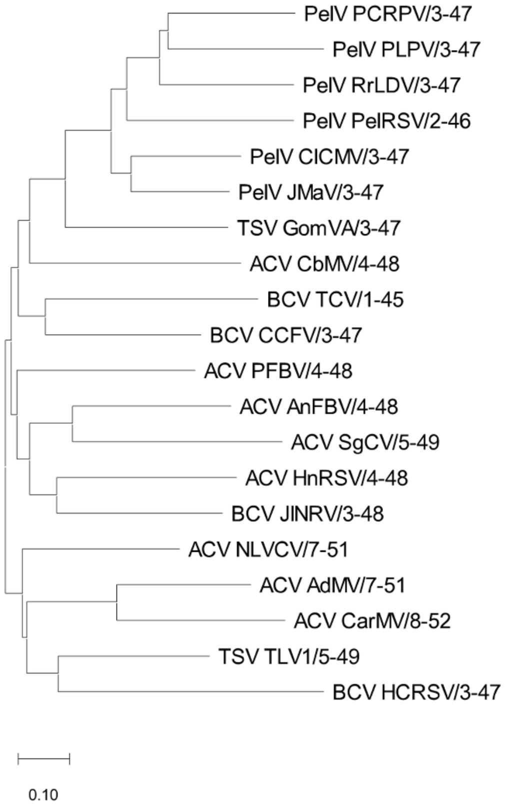
PelV PCRPV/3-47
PelV PLPV/3-47
PelV RrLDV/3-47
PelV PelRSV/2-46
PelV ClCMV/3-47
PelV JMaV/3-47
TSV GomVA/3-47
ACV CbMV/4-48
BCV TCV/1-45
BCV CCFV/3-47
ACV PFBV/4-48
ACV AnFBV/4-48
ACV SgCV/5-49
ACV HnRSV/4-48
BCV JINRV/3-48
ACV NLVCV/7-51
ACV AdMV/7-51
ACV CarMV/8-52
TSV TLV1/5-49
BCV HCRSV/3-47
0.10

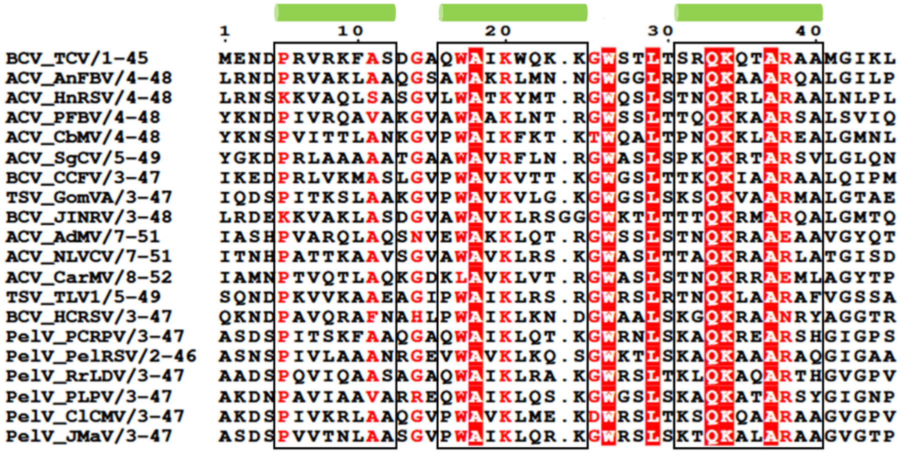
1
10
20
30
40
BCV_TCV/1-45
ACV_AnFBV/4-48
ACV_HnRSV/4-48
ACV_PFBV/4-48
ACV_CbMV/4-48
ACV_SgCV/5-49
BCV_CCFV/3-47
TSV_GomVA/3-47
BCV_JINRV/3-48
ACV_AdMV/7-51
ACV_NLVCV/7-51
ACV_CarMV/8-52
TSV_TLV1/5-49
BCV_HCRSV/3-47
PelV_PCRPV/3-47
PelV_PelRSV/2-46
PelV_RrLDV/3-47
PelV_PLPV/3-47
PelV_ClCMV/3-47
PelV_JMaV/3-47

**Supplementary Figure S13. Structural and sequence alignment of the R-domains of Cluster 1.** (*left*) Dendrogram generated using MEGA 11 with the neighbor-joining algorithm (Poisson model) and 10,000 bootstrap replicates. (*right*) Structure-based sequence alignment of AlphaFold3 models generated in MOE (Molecular Operating Environment); green cylinders indicate predicted α-helices, additionally highlighted by black rectangles within the alignment. For the structural superposition of Cluster 1 models see **Supplementary Fig. 12**.

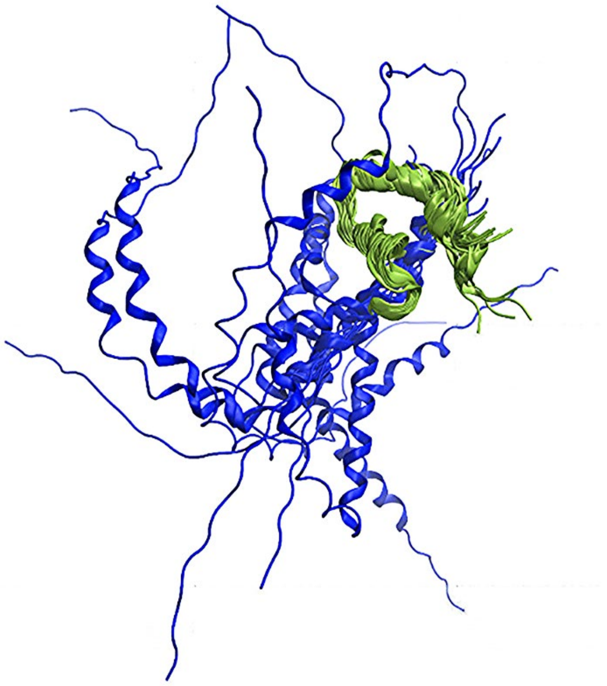

**Supplementary Figure S14. Structural superposition of the N-terminal part of the AlphaFold3 models of *Tombusviridae* capsid proteins (residues 1-45 according to TCV P38).** Structure and sequence-based alignment of multiple *Tombusviridae* family proteins using MOE, based on AlphaFold3-predicted models. The superposition focuses on the N-terminal region of the TCV P38 reference protein (residues 1-45). Models shown in green form a structurally homogenous cluster with pronounced secondary structure (three a-helices), while blue models are coil-dominated and structurally more diverse. This dichotomy corresponds to the clustering observed in the phylogenetic analyses (see **Supplementary Fig. S12**).

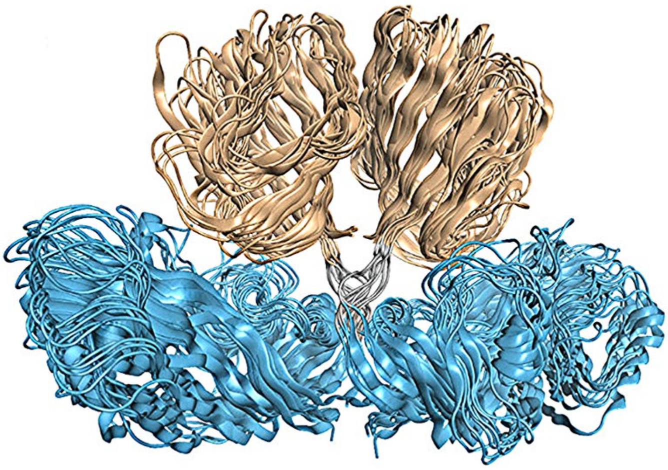

**Supplementary Figure S15**. **Representative dimer models per genus.** Shown is one AlphaFold3-generated dimer model per genus (BCV: TCV, ACV: CarMV, PelV: ClCMV, TSV: GomVA (Cluster 1). AurV: CLSV, DiaV: CRSV, GCV: MNSV, TBV: TBSV, MCV: FNSV (Cluster 2); for abbreviations see **Table S2**). All models exhibit the same relative orientation of the monomers, indicating a conserved dimer topology.
